# Mesaconate-Based Polyester Nanoparticles Reveal How Metabolite Isomerism Shapes Polymer Behavior, Macrophage Reprogramming, and Systemic Immunomodulation

**DOI:** 10.64898/2026.10.22.760948

**Authors:** Md Mohosin Rana, Andrea L. Cottingham, Sara Kolhatkar, Fanny Xu, Alireza Hassani Najafabadi, Ryan M. Pearson

## Abstract

Metabolite-based polymer nanoparticles can transform endogenous immunometabolites into intrinsically bioactive materials, yet most platforms have focused on itaconate or soluble metabolite analogs. Mesaconate is a downstream itaconate-derived immunometabolite with distinct metabolic activity, but its potential as an engineerable material building block remains largely unexplored. Here, mesaconic acid was used as a monomer to construct a degradable, cargo-free polyester nanoparticle platform. Poly(itaconate-co-decanediol) (pITA) and poly(mesaconate-co-decanediol) (pMSA) were synthesized using 1,10-decanediol and formulated into nanoparticles. Varying pMSA synthesis conditions generated distinct molecular-weight development, thermal organization, and degradation-associated behavior. Despite comparable particle size, surface charge, and morphology, pITA- and pMSA-derived nanoparticles produced activation-state-dependent macrophage immunomodulation. Under inflammatory stimulation, both nanoparticles modulated activation-associated surface molecules, whereas their transcriptomic responses diverged markedly. pITA-NPs preferentially engaged redox-, metabolic-, and cellular stress-associated programs, while pMSA-NPs exhibited a distinct lysosome-associated, inflammatory, and remodeling transcriptional profile and remained distinguishable from soluble 4-octyl itaconate. In LPS-induced endotoxemia, pMSA-NP treatment broadly modulated systemic cytokine responses, including a pronounced reduction in circulating IL-6, and improved seven-day survival relative to vehicle control. Together, these findings establish mesaconate as a synthesis-tunable polymeric immunometabolite and highlight metabolite-isomer identity and nanoparticle material state as complementary design parameters for cargo-free immunometabolic biomaterials.

## 1. Background

Endogenous immunometabolites are increasingly recognized as powerful building blocks for engineering intrinsically bioactive materials that interface with immune cell metabolism and function to shape therapeutic responses. The activation state of innate immune cells such as macrophages, have been shown to be tightly coupled to mitochondrial activity, glycolysis, redox balance, and cytokine signaling.^1–3^ In response to pathogens, pathogen- or damage-associated molecular patterns, cytokines, or biomaterial-derived cues, macrophages coordinately adapt their metabolic and signaling pathways to initiate processes involved in host defense, tissue repair, and immune resolution.^2,3^ However, when these pathways become dysregulated, as in inflammatory or other immune-mediated conditions, alterations in macrophage metabolism and signaling can reinforce one another, driving persistent immune activation and other systemic pathophysiological consequences.^1,3,4^ Thus, materials designed to reprogram macrophage immunometabolism may provide a means to restore a more balanced immune response and promote resolution and disease modification.^3,4^

Itaconate is a central macrophage-associated immunometabolite produced during inflammatory activation through remodeling of the mitochondrial tricarboxylic acid (TCA) cycle.^5,6^ In activated macrophages, aconitate decarboxylase 1 (ACOD1), also known as immune-responsive gene 1 (IRG1), converts the TCA-cycle intermediate *cis*-aconitate to itaconate.^5–7^ Itaconate has been shown to dampen macrophage inflammatory responses through mechanisms involving succinate dehydrogenase (SDH) inhibition, altered mitochondrial function and redox balance, glycolytic remodeling, inflammasome regulation, and modulation of inflammatory cytokine production.^6–9^ Its immunomodulatory properties have motivated the development of itaconate derivatives, including dimethyl itaconate (DMI) and 4-octyl itaconate (4OI), as pharmacological tools for targeting macrophage immunometabolism.^8,9^ However, these derivatives do not fully recapitulate endogenous itaconate biology and can introduce pharmacology, dose, solubility, formulation, and off-target toxicity challenges.^9^

Mesaconate, another macrophage-associated immunometabolite, provides a mechanistically distinct extension of the ACOD1-itaconate axis. In inflammatory macrophages, mesaconate is synthesized downstream of itaconate through an itaconate-dependent catabolic process involving succinate-CoA ligase GDP/ADP-forming subunit alpha (SUCLG1)- and AU RNA Binding Methylglutaconyl-CoA Hydratase (AUH).^10^ Although structurally related to itaconate, mesaconate is a distinct C5 dicarboxylate isomer with altered alkene placement and biological activity.^10^ Soluble itaconate and mesaconate both dampen glycolytic activity, whereas only itaconate strongly suppresses TCA-cycle activity, cellular respiration, and SDH function.^10^ Despite these emerging observations, the immunometabolic functions of mesaconate remain far less defined than those of itaconate. This comparatively selective yet incompletely defined bioactivity profile makes mesaconate an attractive building block for developing materials that regulate inflammatory function through immunometabolic mechanisms that are distinct from, but not yet as well characterized as, those of itaconate.

A major challenge in translating endogenous dicarboxylate metabolites into immunomodulatory therapies is intracellular delivery. Many natural metabolites are highly polar and have limited cell-membrane permeability, which can require high extracellular concentrations to achieve intracellular activity.^8,9,11^ Cell-permeable surrogates such as DMI and 4OI partially address this uptake limitation, but their pharmacology differs from the parent metabolite and their use may require repeated high-dose exposure or formulation conditions that are not ideal for translational applications.^8^ Metabolite-based biodegradable polymers provide an alternative strategy, where the endogenous metabolites can be directly incorporated into the polymer backbone, enabling formulation into particles that can be readily internalized by phagocytic immune cells and subsequently degraded intracellularly to release corresponding small-molecule metabolites.^12,13^ Several endogenous metabolites, including succinate, α-ketoglutarate, and itaconate, have been successfully developed into degradable polymer systems, including microparticles and scaffolds for applications spanning cancer immunotherapy, autoimmunity, metabolic disease, and tissue regeneration.^13–18^ Because these metabolites exert distinct and sometimes opposing effects on immune cell function, metabolite identity itself provides a tunable mechanism for directing immunomodulation. Within this framework, itaconate-based polymers have established an important precedent in this area, with recent studies using micro-/nanoparticle systems leveraging enhanced cell uptake and intracellular degradation to modulate macrophage metabolism and inflammatory responses.^13,16,19–21^

Evaluating mesaconate creates an opportunity to extend this concept beyond enhancing metabolite delivery and examine how subtle differences in metabolite structure, when integrated into biomaterials, translate into distinct material properties and biological responses. As a downstream metabolite and structural isomer of itaconate, the different alkene placement of mesaconate may alter how the resulting polymer chains grow, organize, and pack, thereby influencing their thermal transitions, degree of molecular organization, and accessibility of the material to degradation under aqueous or acidic physiological conditions.^3,4,22^ These structural differences may subsequently influence intracellular processing and, ultimately, immune cell transcriptional responses and therapeutic activity.

This relationship is particularly important for cargo-free nanoparticles, where material composition and nanostructure can themselves contribute to multimodal biological activity.^23–26^ Thus, incorporation of itaconate or mesaconate into the polymer backbone changes the context in which these metabolites are presented and delivered to cells, such that their intrinsic bioactivity becomes coupled to the structure and physical state of the resulting material.^27^ In unsaturated polyesters, the position and geometry of the C=C bond can influence amorphous-to-crystalline transitions, chain packing, and thermal transitions,^28^ providing a possible direct mechanism through which metabolite isomerism may alter polymer and nanoparticle organization. Polymer synthesis conditions and nanoparticle formulation add further control over these material properties and, consequently, the biological response. Mesaconate-based nanoparticles would therefore not be expected to simply reproduce the same effects of soluble analogs; rather, their activity may emerge from the interaction between metabolite-derived backbone chemistry, material organization, degradation, and intracellular processing.

Here, we used the isomeric relationship between itaconate and mesaconate to design and mechanistically investigate metabolite-based polyester nanoparticles for macrophage immunomodulation and *in vivo* protection against severe systemic inflammation. The exo-methylene group of itaconate compared to the internal methyl-substituted alkene of mesaconate provides a defined structural difference through which to examine how metabolite identity influences polymer and nanoparticle behavior. We hypothesized that metabolite isomerism would influence polyester synthesis and chain packing, which in turn would shape nanoparticle organization, degradation-associated processing, and macrophage immunomodulatory activity. To test this, we synthesized poly(itaconate-co-decanediol) (pITA_C10_) and poly(mesaconate-co-decanediol) (pMSA_C10_) using 1,10-decanediol (C10) as a hydrophobic linker. For pMSA_C10_, we systematically varied synthesis temperature and inhibitor use to generate a focused formulation series, allowing us to determine how reaction conditions affect polymer organization and nanoparticle bioactivity. The resulting polyesters were formulated into cargo-free nanoparticles and evaluated to define structure-property-bioactivity relationships across polymer synthesis, nanoparticle organization, and macrophage immunomodulation. Polymer and nanoparticle characterization defined relationships among polymer chain growth kinetics, thermal organization, nanoparticle structure, and degradation behavior under lysosome-mimicking acidic conditions. Macrophage phenotyping and RNA sequencing were then used to characterize distinct transcriptional responses of immunometabolic reprogramming elicited by pITA- and pMSA-based nanoparticles and to resolve their major biological differences. Finally, biodistribution, histology, blood chemistry, plasma cytokine profiling, and survival studies in LPS-induced endotoxemia were used to assess systemic distribution, toxicity, immunomodulatory activity, and therapeutic efficacy. Together, this study establishes mesaconate-based polymers and nanoparticles as tunable immunometabolite-based biomaterials, while evaluating metabolite isomerism as a broader design strategy for engineering intrinsically bioactive materials with programmable effects on macrophage immunometabolism and systemic inflammatory responses.

## 2. Results and Discussion

### 2.1 Metabolite-isomer-guided design of itaconate- and mesaconate-based polyesters

To experimentally evaluate metabolite isomerism as a polyester design variable, metabolite-based polymers were synthesized by melt polycondensation of itaconate (ITA) or mesaconate (MSA) with 1,10-decanediol in the presence of stannous chloride (SnCl_2_), with or without 4-methoxyphenol (MeHQ), yielding poly(itaconate-co-decanediol) (pITA_C10_) and three poly(mesaconate-co-decanediol) (pMSA_C10_) polymers, respectively (**Figure 1A,B; Table S1**).

**Figure 1.**
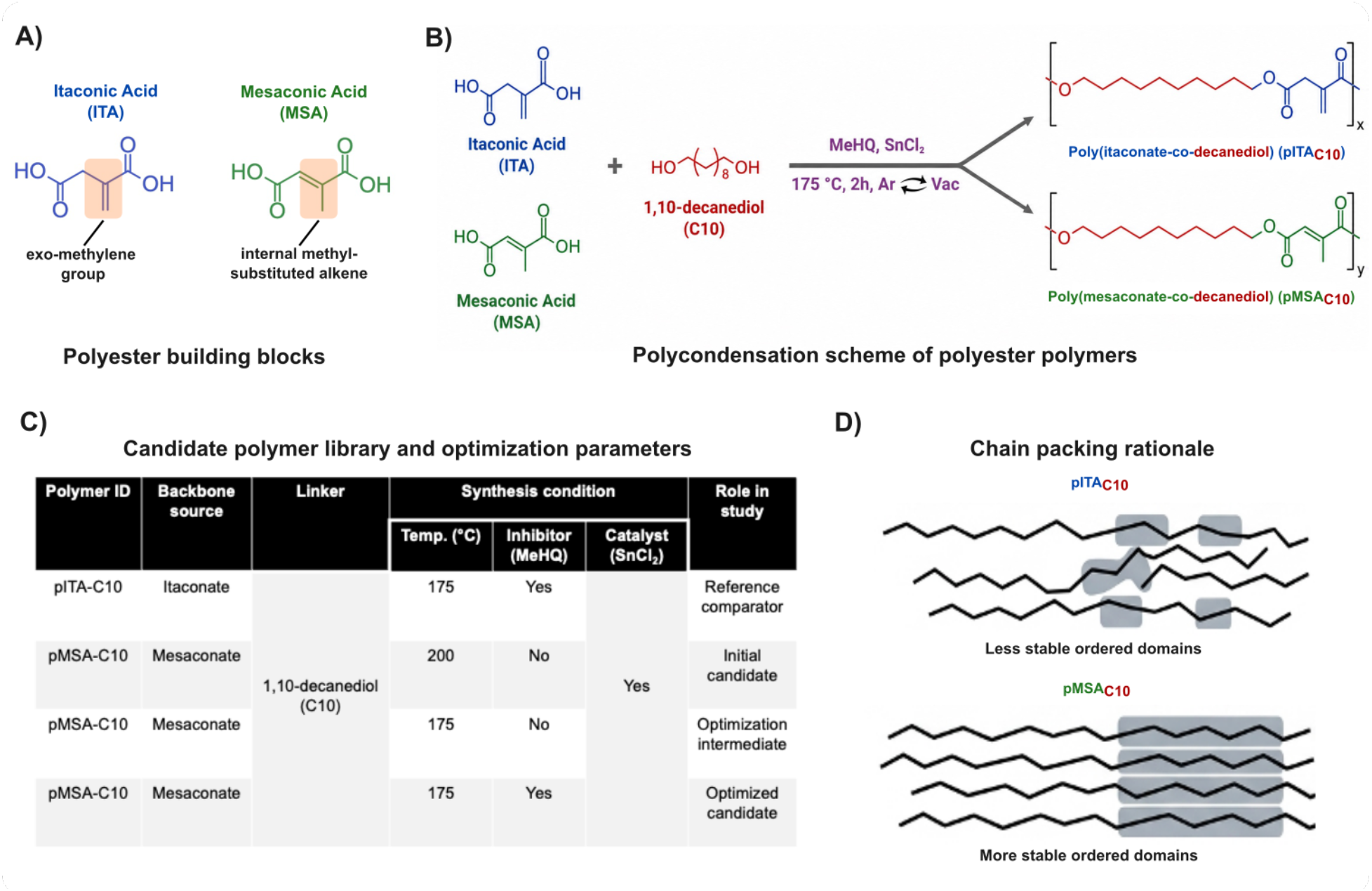
Design and synthesis of metabolite-based polyesters. **(A)** Chemical structures of itaconic acid (ITA) and mesaconic acid (MSA), highlighting the backbone-geometry difference between the exo-methylene group of ITA and the internal methyl-substituted alkene of MSA. **(B)** Schematic representation of polyester synthesis using ITA or MSA with 1,10-decanediol to generate poly(itaconate-co-decanediol) (pITA_C10_) and poly(mesaconate-co-decanediol) (pMSA_C10_), respectively. **(C)** Overview of pMSA_C10_ synthesis conditions used to generate the formulation series. **(D)** Proposed design hypothesis showing that metabolite-isomer backbone geometry and synthesis condition regulate polymer chain organization, nanoparticle accessibility, and macrophage-directed immunomodulatory activity.

pITA_C10_ was synthesized at 175°C in the presence of MeHQ using our previously established melt-polycondensation approach (noted as pITA_C10_^175^ ^°C*^),^21^ which directly incorporates ITA into the polyester backbone rather than relying on DMI as the esterified monomer precursor used in prior pITA polymer syntheses.^16,29–31^ To generate pMSA_C10_ polymers, reaction temperature and inhibitor use were varied resulting in three formulations: (1) pMSA_C10_^200^ ^°C^ synthesized at 200 °C without MeHQ; (2) pMSA_C10_^175^ ^°C^ synthesized at 175 °C without MeHQ; (3) pMSA_C10_^175^ ^°C*^ synthesized at 175 °C with MeHQ (**Figure 1C**). Based on the different alkene substitution patterns of the two monomers, we hypothesized that the pendant exo-methylene group of pITA_C10_ would introduce greater local backbone asymmetry, whereas the internal methyl-substituted alkene of pMSA_C10_ would favor a more conformationally constrained repeat-unit geometry (**Figure 1D**). The resulting polymer library enabled direct evaluation of how metabolite-isomer structure and pMSA synthesis conditions influence polymerization and physicochemical properties.

### 2.2 Polymer reaction kinetics and physicochemical characterization

We next performed extensive physicochemical characterization of the pITA_C10_ and pMSA_C10_ polymers, which revealed clear differences despite their close structural relationship. Gel permeation chromatography (GPC) analysis revealed metabolite- and synthesis condition-dependent differences in molecular-weight distributions across the polymer library (**Figure 2A**). pITA_C10_^175^ ^°C*^ yielded the highest molecular weight among the tested polymers, with an M_n_ of approximately 27.92 ± 2.31 kDa, with a dispersity (Đ) of 2.27 ± 0.31 (**Figure 2B**). In contrast, pMSA_C10_ polymers exhibited lower M_n_ values ranging from 8.54 ± 0.65 to 9.82 ± 0.14 kDa, with corresponding Đ values ranging from 1.57 ± 0.21 to 1.90 ± 0.20, depending on synthesis condition (**Table S2**). The M_n_ achieved for pITA_C10_^175^ ^°C*^ exceeds values reported for several ITA-based polyester systems.^16,32,33^ For example, a recent Davenport-group poly(dodecyl itaconate) synthesized from DMI and 1,12-dodecanediol exhibited an M_n_ of 10.86 ± 1.19 kDa with a Đ of 2.26 ± 0.15.^16^ Notably, pITA_C10_^175^ ^°C*^ reached an approximately 2.6-fold higher M_n_ while maintaining a comparable Đ, indicating that the higher molecular weight was not accompanied by substantial broadening of the molecular-weight distribution. A clear distinction was also observed between the pITA and pMSA polymer systems, with pITA_C10_ exhibiting substantially greater molecular weight than all pMSA_C10_ formulations. Despite variation in synthesis temperature and inhibitor use, the pMSA_C10_ polymers reached a relatively narrow M_n_ range. Because these polymers form through step-growth polycondensation, differences in effective functional-group conversion or competing reactions could produce substantial differences in final molecular weight. Previous studies of itaconate-based polyester synthesis have demonstrated that catalyst identity, reaction conditions, and alkene-associated side reactions can influence esterification behavior and molecular-weight development.^32,34–36^ The distinct exo-methylene structure of ITA and internal methyl-substituted alkene of MSA create different steric and electronic environments around the unsaturated dicarboxylate functionality, which may influence carboxyl-group reactivity, catalyst interactions, or competing reactions during polycondensation. Together, these structural differences provide a plausible chemical basis for the distinct molecular-weight behavior observed between the pITA and pMSA systems.

**Figure 2.**
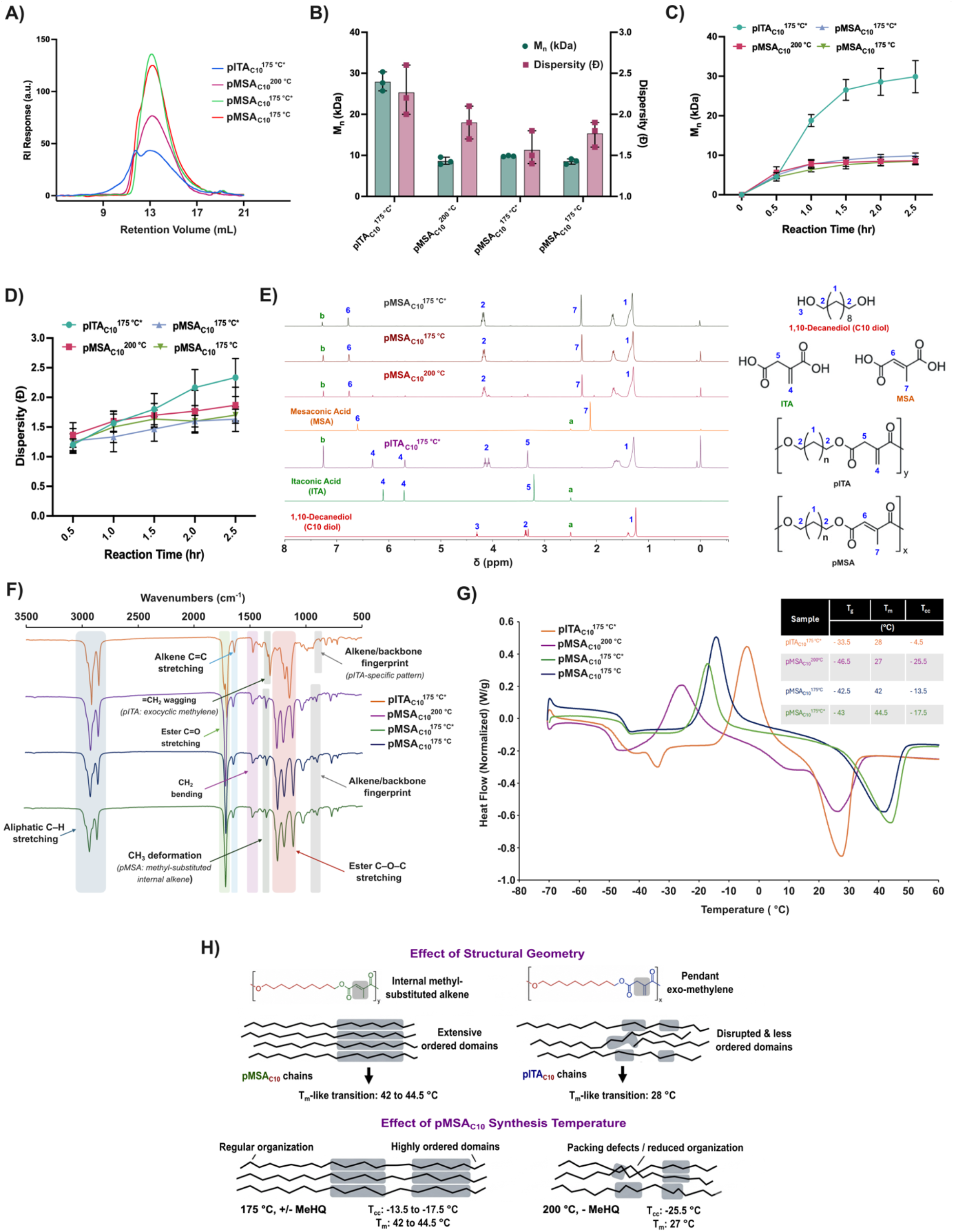
Polymer synthesis optimization and physicochemical characterization. **(A)** Representative GPC traces of pITA_C10_ and pMSA_C10_ formulations showing formulation-dependent molecular weight distributions. **(B)** Quantification of number-average molecular weight (Mn), weight-average molecular weight (Mw), and dispersity (Đ) for pITA_C10_ and pMSA_C10_ formulations. **(C)** Reaction-time-dependent molecular weight development of pITA_C10_ and pMSA_C10_ formulations during polycondensation. **(D)** Corresponding dispersity changes over the reaction period. **(E)** Representative ^1^H NMR spectra confirming successful polyester formation and retention of ITA- or MSA-derived backbone-specific signals. Signals labeled a and b denote the residual DMSO and CHCl_3_ (from CDCl₃) solvent peaks used for NMR analysis. **(F)** FTIR spectra showing ester bond formation and backbone-associated chemical features of pITA_C10_ and pMSA_C10_. **(G)** DSC thermograms comparing thermal transitions of pITA_C10_ and pMSA_C10_ formulations. **(H)** Summary of thermal transition temperatures and enthalpy values, demonstrating synthesis-dependent thermal organization in the pMSA series. Data are presented as mean ± SD.

GPC was next used to assess molecular-weight development over the 2.5 h polymerization period (**Figure 2C,D**). pITA_C10_ exhibited a pronounced increase in M_n_, reaching 29.89 ± 4.07 kDa at 2.5 h, whereas the pMSA_C10_ formulations exhibited more modest increases, reaching M_n_ values of 8.51–9.86 kDa over the same period. Dispersity (Đ) increased from approximately 1.2 to 2.3 for pITA_C10_ and from approximately 1.2–1.4 to 1.6–1.9 across the pMSA_C10_ formulations, indicating differences in molecular-weight development and distribution breadth between the two metabolite-derived polymer systems (**Table S3**).

^1^H NMR and FTIR spectroscopy were used to confirm polyester formation and retention of metabolite-specific structural features (**Figure 2E,F; Figure S1-S4**). For pITA_C10_, characteristic exo-methylene proton resonances were observed at approximately 6.31 and 5.69 ppm, whereas pMSA_C10_ polymers displayed an internal alkene resonance at 6.75-6.76 ppm and the corresponding vinylic methyl signal near 2.26-2.28 ppm. These characteristic resonances were consistently observed across all three pMSA_C10_ formulations, supporting retention of the mesaconate-derived unsaturated structural motif under the investigated synthesis conditions. Across both polymer families, C10-specific signals were observed, including ester-adjacent -OCH_2_-resonances near 4.13-4.19 ppm, methylene signals near 1.60-1.70 ppm, and the internal aliphatic methylene protons near 1.29 ppm. FTIR analysis provided complementary evidence of polyester formation, with aliphatic CH_2_ stretching bands near 2920 and 2850 cm^-1^, ester C=O stretching near 1720-1735 cm^-1^, and ester C-O-C bands in the 1230-1160 cm^-1^ region, consistent with polyester formation.^34,35,37,38^ Bands near 1630-1645 cm^-1^ supported retention of the unsaturated ITA and MSA features.^35,37,38^ pITA_C10_ displayed band stretching in the 890-910 cm^-1^ region, consistent with the exo-methylene group. In contrast, pMSA_C10_ displayed a band near 1370-1380 cm^-1^ attributable to CH_3_ deformation. Because the FTIR features assigned to MSA are not uniquely confirmatory of its structure, they were interpreted in conjunction with the corresponding ^1^H NMR spectra. Collectively, the ^1^H NMR and FTIR results confirmed polyester formation, while supporting retention of the characteristic ITA- and MSA-derived unsaturated structural features across the polymer library.

To further evaluate whether metabolite isomerism influenced polymer chain organization, differential scanning calorimetry (DSC) was used to assess thermal transitions across the pITA and pMSA polymers. pITA_C10_^175^ ^°C*^ showed a melting transition (T_m_) at approximately 28 °C, whereas pMSA_C10_ formulations synthesized at 175 °C showed higher T_m_ values between 42-44.5 °C (**Figure 2G, H**), indicating more thermally stable ordered domains. These differences are consistent with reports that C=C geometry can influence chain organization and crystallization in unsaturated polymers.^22,27^ In the present system, the pendant exo-methylene geometry of pITA_C10_ may disrupt regular chain association relative to the more conformationally constrained internal alkene of pMSA_C10_.

Within the pMSA_C10_ series, the synthesis condition also influenced thermal behavior despite broadly comparable molecular weights. pMSA_C10_^200^ ^°C^ showed a substantially lower T_m_ near 27 °C and a cold-crystallization (T_cc_) feature near -25.5 °C, whereas pMSA_C10_^175^ ^°C^ and pMSA_C10_^175^ ^°C*^ showed higher T_m_ near 42 and 44.5 °C, respectively, with T_cc_ near -13.5 and -17.5 °C. Notably, the pMSA_C10_ formulations retained their characteristic mesaconate-derived unsaturated structural features while exhibiting substantially different thermal responses. The lower melting transition and altered cold-crystallization behavior of the 200 °C formulation is consistent with less stable ordered domains, potentially reflecting differences in chain regularity or microstructural heterogeneity introduced during synthesis. Previous studies have demonstrated that reaction conditions and competing alkene-associated reactions can influence unsaturated polyester formation.^32,35–37^ Although the precise molecular origins of the observed thermal differences remain unresolved, these findings demonstrate that synthesis conditions provide an additional means of modulating the thermal organization of mesaconate-derived polyesters.

Together, these results demonstrate that metabolite-isomer structure and synthesis conditions influence polyester chain growth and thermal organization. While pITA_C10_ achieved higher molecular weight with a comparatively low melting transition, pMSA_C10_ exhibited synthesis-dependent differences in ordered-domain stability, establishing distinct material states for subsequent nanoparticle formulation and biological evaluation.

### 2.3 Nanoparticle formulation and physicochemical properties

The synthesized polyesters were formulated into cargo-free nanoparticles using an oil-in-water emulsion-solvent evaporation method (**Figure 3A**). This cargo-free design enabled evaluation of the metabolite-based polymer itself as the bioactive material without contributions from an encapsulated therapeutic cargo. DLS analysis showed that all formulations produced submicron nanoparticles with broadly comparable hydrodynamic diameters, approximately within a 500–560 nm range (**Figure 3B**). The formulations also showed relatively low polydispersity indices, generally below 0.2, indicating comparatively narrow hydrodynamic size distributions (**Figure 3C**). Zeta-potential values were consistently negative, ranging from approximately −25 to −30 mV (**Figure 3D**). Thus, despite differences in polymer composition and synthesis conditions, the formulations exhibited broadly comparable initial size distributions and surface charge.

**Figure 3.**
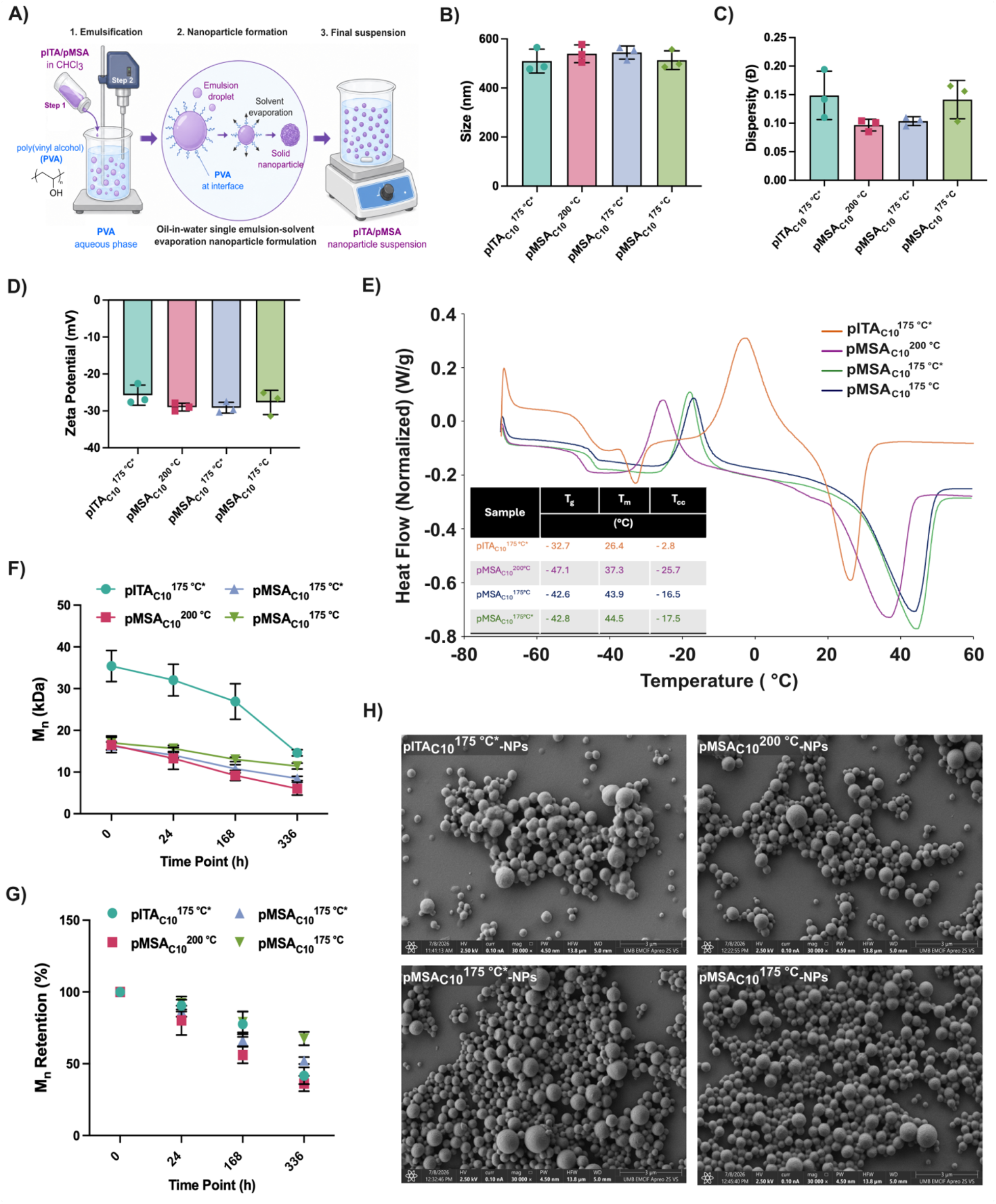
Nanoparticle formulation, nanoparticle-state organization, and acidic degradation-associated behavior of pITA_C10_ and pMSA_C10_ nanoparticles. **(A)** Schematic illustration of cargo-free nanoparticle fabrication by an oil-in-water single-emulsion solvent evaporation method. pITA_C10_ or pMSA_C10_ polymers dissolved in CHCl_3_ were emulsified into an aqueous PVA phase, followed by solvent evaporation to generate polymer nanoparticles. **(B)** Hydrodynamic diameter of pITA_C10_- and pMSA_C10_-based nanoparticles measured by dynamic light scattering. **(C)** Dispersity index of the nanoparticle formulations. **(D)** Zeta potential of the nanoparticle formulations, showing negatively charged surfaces across all groups. **(E)** DSC thermograms of lyophilized nanoparticles with corresponding thermal transition parameters, including T_g_, T_m_, and T_cc_ values, indicating that synthesis-dependent thermal behavior was largely retained after nanoparticle formulation. **(F)** GPC-derived number-average molecular weight (M_n_) of the recovered polymer fraction after acidic incubation in pH 5.2 acetate buffer at 37 °C. **(G)** Corresponding M_n_ retention (%) during acidic incubation, showing formulation-dependent molecular-weight reduction under macrophage-relevant acidic conditions. **(H)** Representative SEM images showing spherical nanoparticle morphology for pITA_C10_ and pMSA_C10_ formulations. Data are presented as mean ± SD where applicable. * denotes inhibitor-containing synthesis condition.

DSC analysis of lyophilized nanoparticles showed that synthesis-dependent thermal organization was largely retained after nanoparticle formulation (**Figure 3E**). pITA_C10_^175^ ^°C*^-NPs showed a melting-like transition (T_m_) near 26.4 °C, whereas pMSA_C10_^175^ ^°C^-NPs and pMSA_C10_^175^ ^°C*^-NPs retained higher T_m_ near 43.9 and 44.8 °C, respectively. In contrast, pMSA_C10_^200^ ^°C^-NPs displayed a T_m_ near 37.3 °C, higher than the corresponding bulk pMSA_C10_^200^ ^°C^ transition but still lower than the 175 °C pMSA nanoparticle formulations. This upward shift may reflect processing-induced chain reorganization and preferential retention of higher-molecular-weight polymer fractions during nanoparticle formation and purification. These data indicate that the synthesis-dependent differences established in the bulk pMSA polymers persist after nanoparticle processing, with pMSA_C10_^200^ ^°C^-NPs retaining lower thermal organization than the two 175 °C formulations. Its transition near physiological temperature may also permit greater chain mobility than the more thermally stable 175 °C pMSA formulations, although the DSC measurements themselves do not directly establish behavior in the cellular environment.^28,39^

We next assessed the time- and formulation-dependent changes in polymer molecular weight after exposure of nanoparticles to acidic buffer representative of the endosome (**Figure 3F,G, Table S4**). Nanoparticles were incubated in pH 5.2 acetate buffer at 37 °C to approximate acidic endosomal/lysosomal-like processing conditions, followed by polymer recovery and GPC analysis. The initial M_n_ values of the recovered nanoparticle-associated polymers were higher than those of the corresponding bulk polymers reported in **Figure 2B**. This difference may reflect molecular-weight fractionation during nanoparticle fabrication and purification, potentially resulting from preferential retention of higher-molecular-weight chains in the recovered nanoparticle fraction. Because the contributions of polymer processing and recovery were not independently resolved, molecular-weight retention was calculated relative to the formulation-specific recovered-polymer M_n_ at 0 h, with the 0 h samples briefly exposed to acidic buffer and immediately washed and recovered to provide a processing-matched baseline. All formulations showed time-dependent decreases in M_n_, consistent with acid-associated polyester chain scission. pITA_C10_^175^ ^°C*^-NPs decreased from an initial recovered M_n_ of 35.40 ± 3.73 kDa to 14.62 ± 0.74 kDa at 336 h, corresponding to 41.7 % Mₙ retention. Among the pMSA-NPs, pMSA_C10_^200^ ^°C^-NP exhibited the greatest molecular-weight reduction, decreasing from 16.49 ± 1.86 to 6.04 ± 1.55 kDa and retaining 36.3 % of their initial M_n_ at 336 h. By comparison, pMSA_C10_^175^ ^°C^-NP and pMSA_C10_^175^ ^°C*^-NP retained 67.5 % and 52.2 %, respectively. These results demonstrate synthesis-dependent differences in the acidic molecular-weight reduction of pMSA-NPs.

Within the pMSA-NPs, acidic molecular-weight reduction corresponded to nanoparticle DSC behavior. pMSA_C10_^200^ ^°C^-NPs, which displayed the lowest melting-like transition among the pMSA formulations near 37.3 °C, also showed the greatest M_n_ reduction, retaining only 36.3 % of the initial M_n_ at 336 h. In contrast, pMSA_C10_^175^ ^°C^-NPs and pMSA_C10_^175^ ^°C*^-NPs retained higher melting-like transitions near 43.9 and 44.8 °C, respectively, and greater molecular-weight retention during acidic incubation. These findings suggest a potential contribution of synthesis-dependent polymer organization to acidic degradation-associated behavior, although the precise molecular origins remain unresolved.^22,27,28^ Although pITA_C10_^175^ ^°C*^-and pMSA_C10_^200^ ^°C^-NPs exhibited comparable final M_n_ retention percentages, their absolute molecular weights remained substantially different, at 14.62 ± 0.74 and 6.04 ± 1.55 kDa, respectively.

DLS analysis under the same acidic incubation conditions showed that molecular-weight reduction and colloidal restructuring were distinct formulation-dependent processes (**Figure S5**). DLS and GPC therefore provide complementary readouts of hydrodynamic particle behavior and polymer-chain molecular weight, respectively.^40,41^ pITA_C10_^175^ ^°C*^-NPs exhibited pronounced hydrodynamic diameter increases, with approximately 60 % increase at 24 h, 180–190 % at 168 h, and 135–145 % at 336 h, consistent with substantial colloidal restructuring, swelling, or aggregation. In contrast, pMSA_C10_^200^ ^°C^-NPs maintained a nearly unchanged hydrodynamic diameter, with size changes close to 0–5 %, despite showing the strongest M_n_ reduction among the pMSA formulations. Thus, substantial molecular-weight reduction in pMSA_C10_^200^ ^°C^-NPs occurred without major changes in hydrodynamic size. The 175 °C pMSA formulations showed only modest colloidal changes, with final diameter increases of approximately 20–30 % at 336 h. Together, these data distinguish polymer-chain molecular-weight reduction from particle-level swelling or aggregation and show that pMSA_C10_^200^ ^°C^-NPs maintained comparatively stable colloidal dimensions despite greater M_n_ loss.

Taken together, the DSC, GPC, and DLS results show that pMSA_C10_^200^ ^°C^-NPs combine lower thermal organization and greater acid-associated molecular-weight reduction with comparatively stable hydrodynamic dimensions. This behavior distinguishes the 200 °C formulation from the more thermally organized 175 °C pMSA nanoparticles and demonstrates that synthesis conditions remain an important determinant of nanoparticle material state after formulation.

SEM imaging confirmed spherical nanoparticle morphology across pITA_C10_ and pMSA_C10_ formulations (**Figure 3H**). Some clustering was observed in the dried-state images, but the images confirmed discrete spherical particles rather than irregular polymer precipitates. Overall, the formulations shared comparable initial colloidal properties and spherical morphology but retained distinct synthesis-dependent thermal and acidic-response profiles. Accordingly, the acidic degradation data should be viewed as a material-level indicator of formulation-dependent acid susceptibility and accessibility when interpreting the subsequent biological studies, rather than as evidence of complete polymer degradation over the timescale of macrophage exposure.

### 2.4 Macrophage bioactivity, viability, and apoptosis

We next evaluated whether pITA- and pMSA-based nanoparticles modulate macrophage activation-associated phenotypes while maintaining cellular compatibility. pITA_C10_^175^ ^°C*^-NPs were used as the pITA benchmark based on our previous optimization of pITA nanoparticle composition and molecular weight, in which the high-molecular-weight C10 formulation exhibited the strongest macrophage phenotypic activity among the evaluated pITA formulations.^21^ Murine bone marrow-derived macrophages were treated with soluble 4OI, pITA_C10_-based nanoparticles, or the pMSA_C10_ nanoparticle formulations under unstimulated or LPS-stimulated (M1) conditions, followed by viability, apoptosis, and flow-cytometric phenotyping after 24 h of incubation (**Figure 4A**; **Figure S6**).

**Figure 4.**
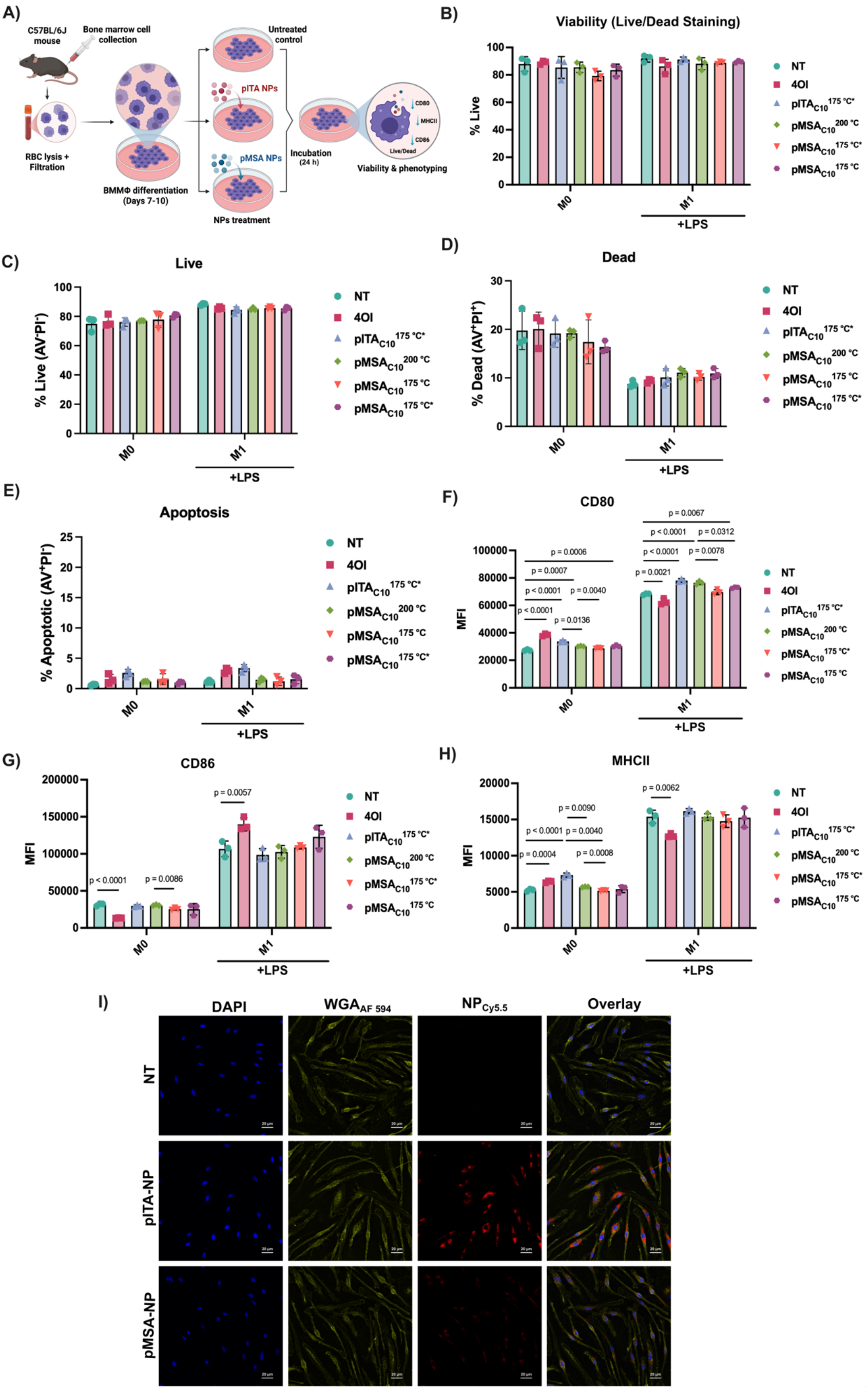
pITA- and pMSA-derived nanoparticles remodel macrophage activation-associated surface markers while preserving viability and limiting apoptotic stress. **(A)** Experimental schematic showing isolation of murine bone marrow cells, differentiation into bone marrow-derived macrophages, treatment with soluble 4OI, pITA_C10_-derived nanoparticles, or pMSA_C10_-derived nanoparticles under unstimulated (M0) or LPS-stimulated (M1-like) conditions, followed by viability, apoptosis, and flow-cytometric phenotyping after 24 h. **(B)** Cell viability determined by Live/Dead staining. **(C)** Annexin V⁻/PI⁻ (AV⁻/PI⁻) live-cell population determined by AV/PI staining. **(D)** AV⁺/PI⁺ dead/late-stage cell population. **(E)** AV⁺/PI⁻ apoptotic-cell population. **(F–H)** Flow-cytometric quantification of macrophage activation-associated markers, including **(F)** CD80, **(G)** CD86, and **(H)** MHCII, reported as median fluorescence intensity (MFI). pITA- and pMSA-derived nanoparticles maintained high viability and low apoptotic fractions while inducing formulation- and activation-state-dependent remodeling of CD80, CD86, and MHCII expression. **(I)** Representative confocal images showing uptake of Cy5.5-labeled pITA-NPs and selected pMSA-NPs by macrophages. DAPI, nuclei; WGA_AF594_, cell membrane; Cy5.5, nanoparticles. Scale bars, 20 µm. Data are presented as mean ± SD, with individual points representing biological replicates. Statistical analyses were performed separately for unstimulated and LPS-stimulated conditions using one-way ANOVA followed by Dunnett’s multiple-comparisons test against the corresponding control. Selected pairwise comparisons between treatment groups were performed using two-sided unpaired t-tests with Welch’s correction. Exact p values are shown where indicated.

Across both unstimulated and LPS-stimulated macrophages, nanoparticle-treated groups maintained high cellular compatibility based on Live/Dead staining (**Figure 4B**). In unstimulated macrophages, viability ranged from approximately 79 % to 88 % across nanoparticle-treated groups, compared with approximately 88 % in untreated cells. Under LPS stimulation, viability remained approximately 90–92 %, compared with approximately 94 % in LPS-treated controls. Consistently, Annexin V/PI analysis showed that most nanoparticle-treated macrophages remained within the AV^-^/PI^-^ live-cell population, with fractions of approximately 76–81 % and 86–88 % under unstimulated and LPS-stimulated conditions, respectively (**Figure 4C**). The AV^+^/PI^+^ population accounted for approximately 16–19 % of unstimulated and 10–11 % of LPS-stimulated nanoparticle-treated macrophages (**Figure 4D**), whereas AV^+^/PI^-^ early-apoptotic cells represented only approximately 3–4 % or less across conditions (**Figure 4E**).^42^ Together, these findings indicate that nanoparticle treatment was not associated with extensive early apoptosis or loss of viability under the tested conditions, supporting subsequent evaluation of macrophage phenotypic responses.

Flow-cytometric analysis revealed formulation- and activation-state-dependent changes in CD80, CD86, and MHCII expression (**Figure 4F-H**). CD80 and CD86 are costimulatory molecules associated with macrophage activation and antigen-presenting function, whereas MHCII is associated with antigen presentation. The observed responses varied across these markers, suggesting selective modulation of activation-associated surface-marker expression rather than a uniform phenotypic shift.

In unstimulated macrophages, pMSA_C10_^200^ ^°C^-NPs increased CD80 expression by approximately 11 %, compared with increases of approximately 24 % and 41 % following pITA_C10_^175^ ^°C*^-NPs and soluble 4OI treatment, respectively (**Figure 4F**). In contrast, CD86 remained largely unchanged following pMSA_C10_^200^ ^°C^-NPs treatment, whereas soluble 4OI reduced CD86 expression by approximately 57 % (**Figure 4G**). MHCII expression increased by approximately 40 % following pITA_C10_^175^ ^°C*^-NP treatment but only approximately 7–10 % following pMSA_C10_^200^ ^°C^-NP treatment (**Figure 4H**). These marker-specific differences indicate that pMSA_C10_^200^ ^°C^-NP formulation generated a surface-marker profile distinct from those observed with pITA_C10_^175^ ^°C^-NPs and soluble 4OI.

Under LPS stimulation, pITA_C10_^175^ ^°C*^-NPs and pMSA_C10_^200^ ^°C^-NPs both enhanced CD80 expression by approximately 15 % and 12 %, respectively, relative to LPS-treated controls (**Figure 4F**). Smaller increases were observed with pMSA_C10_^175^ ^°C*^-NPs (∼3 %) and pMSA_C10_^175^ ^°C^-NPs (∼6–7 %). CD86 expression remained close to the LPS control following pITA_C10_^175^ ^°C*^-NP and pMSA_C10_^200^ ^°C^-NP treatment, whereas soluble 4OI treatment increased CD86 expression by approximately 31 % (**Figure 4G**). In contrast, MHCII expression changed only modestly following pITA_C10_^175^ ^°C*^-NP treatment (approximately 4–5 %) and remained largely unchanged following pMSA_C10_^200^ ^°C^-NP treatment (∼0–2 % variation) (**Figure 4H**). Together, these results demonstrate that the metabolite-based nanoparticles produced marker-specific modulation rather than uniformly activating phenotypic responses under inflammatory stimulation.

The pMSA nanoparticle formulations also exhibited synthesis condition-associated differences in macrophage bioactivity. Among the pMSA formulations, pMSA_C10_^200^ ^°C^-NPs produced the clearest CD80-associated response under LPS stimulation, whereas the two 175 °C formulations elicited smaller changes. When considered alongside the materials characterization in **Figures 2-3**, the 200 °C formulation combined lower nanoparticle thermal organization and greater acid-associated M_n_ reduction with the most distinct CD80 response among the pMSA formulations. On this basis, pMSA_C10_^200^ ^°C^-NPs were selected for subsequent transcriptomic and *in vivo* evaluation.

Confocal fluorescence microscopy further showed nanoparticle-associated fluorescence within macrophages following exposure to pITA-NPs and the selected pMSA_C10_^200^ ^°C^-NPs, consistent with cellular uptake of both formulations (**Figure 4I**). Although the pITA- and pMSA-NPs exhibited broadly comparable initial hydrodynamic size, size dispersity, surface charge, and spherical morphology, the formulations differed in molecular weight and synthesis history. Therefore, the observed biological differences should not be attributed solely to metabolite isomerism but rather interpreted in the context of the distinct physicochemical states of the two nanoparticle platforms. Together with the synthesis-dependent thermal and acidic-response profiles established in **Figure 3**, these findings support further investigation of polymer chemistry and material organization as potential contributors to macrophage phenotypic modulation. However, intracellular trafficking and intracellular polymer degradation will be required to establish the mechanisms connecting these material properties to macrophage bioactivity.^43–47^ Overall, pITA- and pMSA-derived nanoparticles preserved macrophage viability while producing formulation- and activation-state-dependent remodeling of CD80, CD86, and MHCII . The selected pMSA_C10_^200^ ^°C^-NP formulation is hereafter referred to as pMSA-NP for subsequent transcriptomic and endotoxemia studies.

### 2.5 RNA-seq reveals distinct macrophage immunometabolic reprogramming by metabolite-based nanoparticles

To determine whether the phenotypic differences observed in **Figure 4** were associated with broader changes in gene expression, genome-wide transcriptional profiling was performed using RNA sequencing (RNA-seq). Unstimulated (M0) and LPS-stimulated bone marrow-derived macrophages were treated with pITA_C10_^175^ ^°C*^-NPs or pMSA_C10_^200^ ^°C^-NPs for 24 h prior to RNA isolation and sequencing (**Figure 5A**). IL-4 stimulated macrophages were used to represent an alternatively activated macrophage control. 4OI was included as a mechanistically distinct control because it is a cell-permeable ITA derivative with established immunoregulatory and redox activity.^7,8^ This comparison allowed us to determine whether various metabolite-based NPs produced transcriptional responses similar to or distinct from those induced by 4OI. Macrophage polarization controls were also included to assess the relationship of these responses to established LPS-associated inflammatory and IL-4-associated alternatively activated transcriptional states.

**Figure 5.**
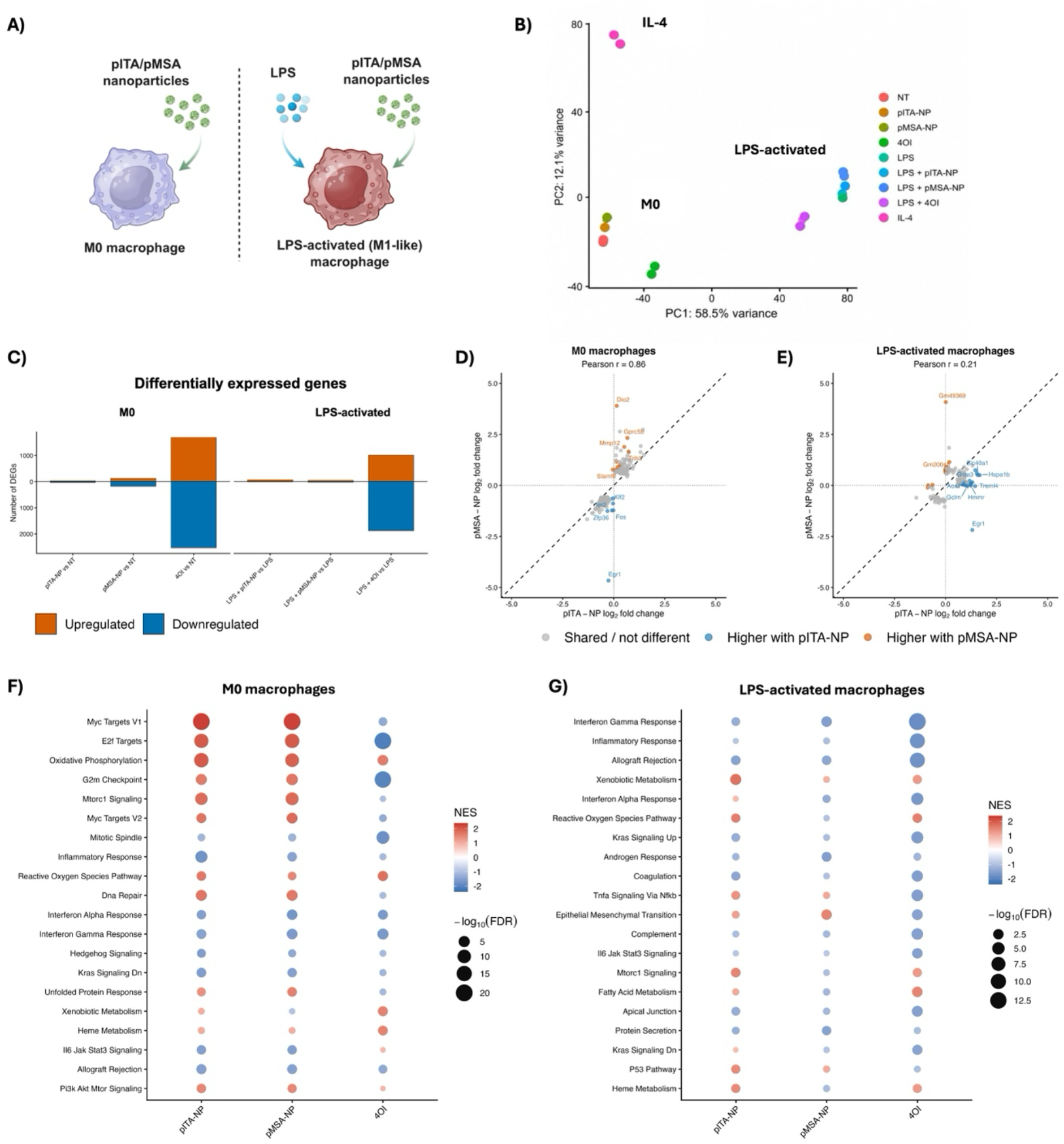
pITA- and pMSA-derived nanoparticles elicit distinct macrophage transcriptional responses. **(A)** Experimental design for RNA-seq analysis of unstimulated (M0) and LPS-activated bone marrow-derived macrophages treated with pITA-NP or pMSA-NP for 24 h. Soluble 4OI and IL-4-treated macrophages were included as controls. **(B)** Principal component analysis (PCA) of global gene-expression profiles across untreated, LPS-activated, IL-4-treated, 4OI-treated, and NP-treated macrophages. **(C)** Number of differentially expressed genes (DEGs) following pITA-NP, pMSA-NP, or 4OI treatment in M0 and LPS-activated macrophages relative to their respective untreated or LPS controls. DEGs were defined by adjusted p < 0.05 and |log_2_ fold change| ≥ 0.58. Correlation analysis comparing pITA-NP- and pMSA-NP-induced gene-expression changes in **(D)** M0 macrophages and **(E)** LPS-activated macrophages. Each point represents an individual gene plotted according to its log_2_ fold change following pITA-NP and pMSA-NP treatment relative to the corresponding control. Gene set enrichment analysis (GSEA) using MSigDB Hallmark gene sets in **(F)** M0 and **(G)** LPS-activated macrophages following pITA-NP, pMSA-NP, or 4OI treatment relative to the corresponding control. Dot color represents the normalized enrichment score (NES), with positive and negative values indicating positive and negative enrichment, respectively, and dot size represents −log_10_(FDR). RNA-seq was performed using n = 3 biological replicates per condition.

Principal component analysis (PCA) showed clear separation of M0, LPS-stimulated, and IL-4-treated macrophages, whereas NP-treated cells remained closer to their corresponding M0 or LPS controls (**Figure 5B**). In contrast, 4OI-treated samples clustered distinctly from both the corresponding controls and NP-treated groups. Consistent with the PCA, 4OI induced a substantially greater number of differentially expressed genes (DEGs; adjusted p < 0.05 and |log_2_FC|≥ 0.58) than either NP formulation (**Figure 5C**; **Table S6**). In M0 macrophages, pITA-NP and pMSA-NP altered 71 and 304 genes, respectively, compared with 4,222 genes following 4OI treatment. Similarly, in LPS-activated macrophages, pITA-NP and pMSA-NP altered 80 and 74 genes, respectively, compared with 2,892 genes following 4OI treatment. Together, these findings indicate that pITA-NP and pMSA-NP produce more focused transcriptional changes, whereas 4OI induces substantially broader global transcriptional reprogramming.

To define shared and treatment-specific transcriptional responses, differential gene-expression and pathway analyses were performed following pITA-NP, pMSA-NP, and 4OI treatment in M0 and LPS-activated macrophages. In M0 macrophages, pITA-NP and pMSA-NP showed substantial overlap in their enriched pathways (**Figure S7**). Both NP formulations positively enriched oxidative phosphorylation and pathways associated with protein synthesis and proteostasis, including ribosomal and proteasomal programs, while showing negative enrichment of cytokine-cytokine receptor interactions, hematopoietic cell lineage, extracellular matrix and cell adhesion, and lysosome-associated pathways.

Visualization of genes associated with inflammatory signaling, immunoregulation, immediate-early signaling, metabolic reprogramming, lysosomal/phagocytic function, and oxidative-stress response, further highlighted both shared and formulation-specific patterns (**Figure S8A**). Closer evaluation of DEGs following treatment with pITA-NP and pMSA-NP highlighted several similarities and differences in the induced transcriptional changes (**Figure S8B,C**). The overall similarity in gene-level responses was supported by a strong correlation between pITA-NP- and pMSA-NP-induced transcriptional changes (Pearson r = 0.86) (**Figure 5D**). Specifically, both formulations increased expression of genes associated with redox regulation and metabolic adaptation, including *Gclm*, *Srxn1*, and *Mthfd2*, while reducing *Ms4a4a*, a macrophage-associated gene implicated in macrophage activation and inflammatory responses. Despite these similarities, pMSA-NP showed higher expression of *Dio2*, a regulator of thyroid hormone activation and thermogenic metabolism, together with stress- and remodeling-associated genes including *Trib3* and *Mmp12*. In contrast, immediate-early and stimulus-responsive genes, including *Egr1*, *Zfp36*, and *Fos*, were expressed at lower levels following pMSA-NP treatment than pITA-NP. 4OI, on the other hand, shared positive enrichment of oxidative phosphorylation and ribosomal programs with both NP formulations but produced a broader and distinct pathway-level response, characterized by enrichment of xenobiotic- and reactive oxygen species-associated metabolism together with negative enrichment of cell-cycle, DNA-replication, and DNA-repair pathways that was not observed with pITA-NP or pMSA-NP (**Figure S9**).

Under LPS stimulation, the transcriptional programs induced by pMSA-NP and pITA-NP diverged most strongly (Pearson r = 0.21) (**Figure 5E**). Despite their differences, both NP formulations showed negative enrichment of immune and trafficking pathways (**Figure S10**), including chemokine signaling and leukocyte transendothelial migration, indicating shared modulation of inflammatory signaling and immune-cell trafficking. The formulations differed more clearly among positively enriched pathways. pITA-NP showed positive enrichment of glutathione metabolism, carbon and xenobiotic metabolism, ferroptosis, and endoplasmic-reticulum protein processing, consistent with a prominent redox, metabolic, and cellular-stress response, whereas pMSA-NP showed positive enrichment only for lysosome biogenesis.

Targeted gene visualization (**Figure S11A**) and DEG analyses (**Figure S11B,C**) further revealed that pITA-NP had higher expression of redox- and stress-associated genes, including *Gclm*, *Gsta3*, and *Hspa1b*, relative to pMSA-NP, consistent with its stronger redox/stress-associated pathway enrichment. In contrast, pMSA-NP showed higher expression of inflammatory and remodeling-associated genes, including *Mmp8*, *Cxcl3*, and *Pf4*, together with lower expression of *Egr1*, *Tmem176b*, and *Slamf9*. Notably, *Gdf15* was among the more strongly induced genes following treatment with both NP formulations. GDF15 has been identified as an inflammation-induced mediator of tissue tolerance that supports survival during acute inflammatory and infectious stress.^48^ Comparison of 4OI and pITA-NP identified several shared metabolic features, including positive enrichment of glutathione, carbon, xenobiotic, and endoplasmic-reticulum-associated pathways, although 4OI produced a substantially broader transcriptional response (**Figure S12A**). For example, 4OI induced higher expression of the NRF2-responsive antioxidant genes *Nqo1* and *Srxn1*, consistent with the established ability of 4OI to activate KEAP1-NRF2 signaling. In contrast, pITA-NP showed higher expression of immune- and chemokine-associated genes, including *Fpr2*, *Ccr5*, and *Ccl12* (**Figure S12B**).

Collectively, pITA-NP and pMSA-NP shared a core immunomodulatory response characterized by suppression of inflammatory signaling and immune-cell trafficking pathways, while differing in the regulation of redox, metabolic, and stress-response programs in a cell activation state-dependent manner. GSEA using the MSigDB Hallmark gene sets further supported this relationship (**Figure 5F,G**). In M0 macrophages, both formulations showed positive enrichment of oxidative phosphorylation and reactive oxygen species-associated programs together with negative enrichment of inflammatory, interferon, and IL-6/JAK/STAT3-associated gene sets, consistent with a coordinated shift in metabolic and inflammatory transcriptional programs. Under LPS stimulation, the responses were more distinct, with pITA-NP retaining stronger enrichment of reactive oxygen species- and metabolism-associated gene sets, whereas pMSA-NP showed a more limited metabolic and redox-associated response. These pathway-level differences were accompanied by distinct regulation of immediate-early, immunoregulatory, and remodeling-associated genes by pMSA-NP. Thus, despite their structural similarity, incorporation of the two metabolite isomers into NPs produced overlapping but distinguishable transcriptional responses that were influenced by macrophage activation state. Importantly, pMSA-NPs did not simply recapitulate the transcriptional profile of either pITA-NPs or soluble 4OI, suggesting that polymer-based MSA provides access to an additional immunometabolic response space beyond established ITA-based approaches. These findings motivated us to next determine whether these formulation-dependent transcriptional differences were associated with distinct systemic biological responses *in vivo*.

### 2.6 In vivo distribution, systemic immunomodulation, and functional validation in LPS-induced endotoxemia

To determine whether the observed *in vitro* biological activity translated into functional benefit *in vivo*, we evaluated the ability of nanoparticle treatment to prevent LPS-induced acute systemic inflammatory stress (**Figure 6A**). Mice received vehicle, pITA-NP, or pMSA-NP before LPS challenge, followed by survival monitoring for seven days. This model was selected as a stringent proof-of-concept model of acute severe systemic inflammatory challenge rather than a disease-specific therapeutic model, although this model has been used previously to evaluate protective responses against cytokine storm.^24,49–51^ Organ-level IVIS imaging of intraperitoneally administered Cy5.5-labeled nanoparticles showed that both pITA-NP and pMSA-NP produced their strongest fluorescence signals in the liver, with significantly greater hepatic fluorescence than the untreated and LPS control groups (p < 0.0001). Lower signals were detected in the lungs, spleen, kidneys, and heart. Representative *ex vivo* organ images corroborated these findings, indicating preferential hepatic accumulation or retention of nanoparticle-associated fluorescence under the experimental conditions (**Figure 6B,C**).

**Figure 6.**
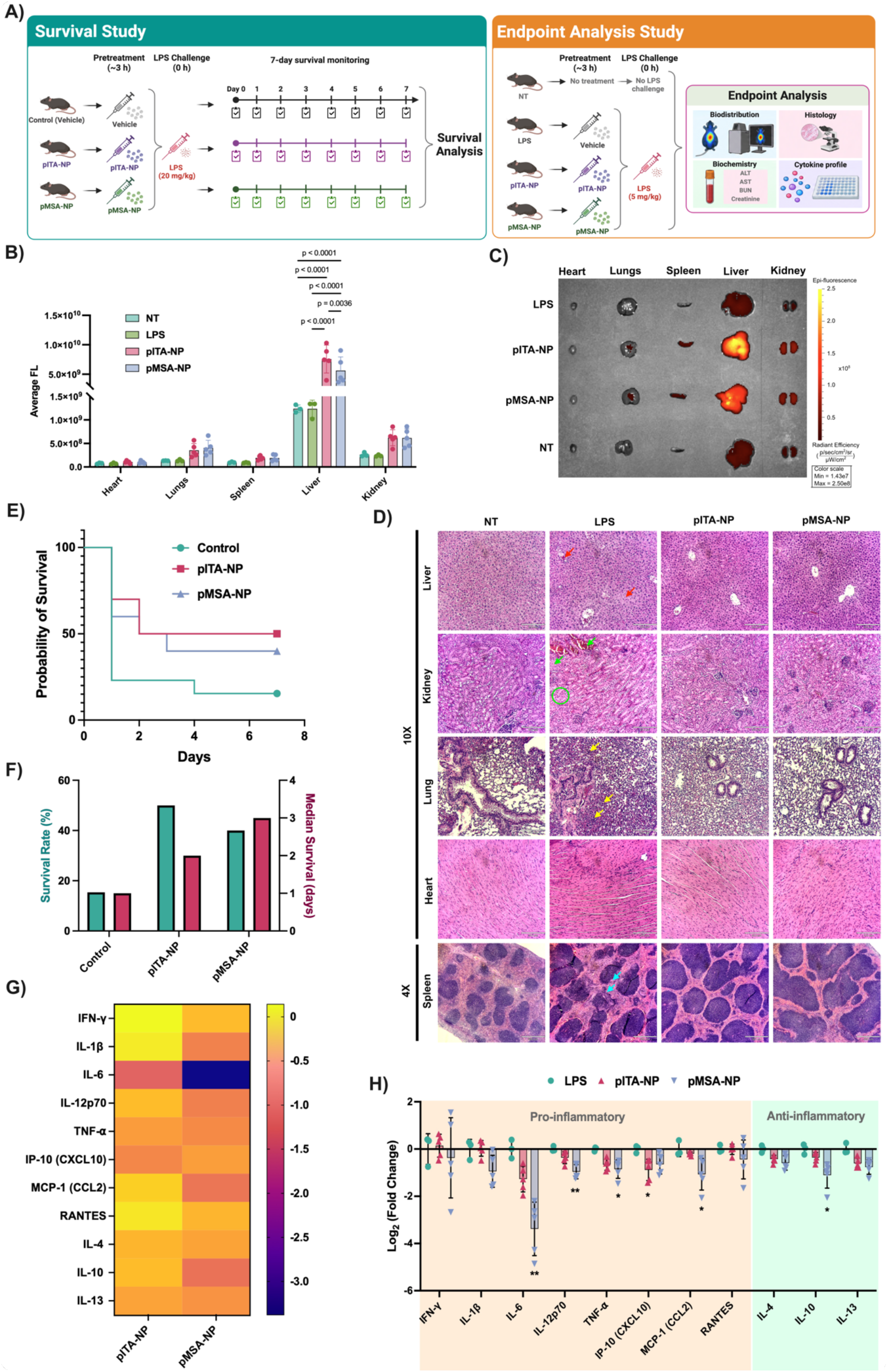
In vivo distribution, systemic cytokine responses, and survival outcomes following metabolite-derived polyester nanoparticle pretreatment in endotoxemia. **(A)** Experimental design of the survival (20 mg kg^-1^ LPS; 7-day monitoring) and endpoint-analysis (5 mg kg^-1^ LPS; biodistribution, histology, blood chemistry, and cytokines) studies. **(B)** Ex vivo fluorescence quantification in major organs, showing predominant hepatic nanoparticle-associated signal. **(C)** Representative fluorescence images of harvested organs. **(D)** Representative H&E-stained sections of liver, kidney, lung, heart, and spleen from NT, LPS, pITA-NP + LPS, and pMSA-NP + LPS groups. Liver, kidney, lung, and heart are shown at 10× (scale bars, 210 µm) and spleen at 4× (520 µm). Arrows/circles indicate representative morphological alterations observed in the LPS group; overall tissue architecture remained identifiable in nanoparticle-treated groups. **(E)** Kaplan–Meier survival curves following LPS challenge in vehicle control (n = 13), pITA-NP (n = 10), and bioactivity-selected pMSA-NP (n = 10) groups. Survival distributions were compared using the log-rank (Mantel–Cox) test with Holm–Šídák correction for multiple comparisons against vehicle control (adjusted P = 0.1204 for pITA-NPs and P = 0.1793 for pMSA-NPs). **(F)** Final survival rates and median survival times for the corresponding groups. **(G)** Heatmap of selected circulating cytokines and chemokines measured by a 26-plex Luminex assay, expressed as mean log₂ fold change relative to LPS control. **(H)** Individual-animal log_2_ fold-change values for the corresponding cytokines, illustrating treatment-associated systemic cytokine responses following nanoparticle pretreatment. Data in **(B)** are presented as mean ± SD with individual values and were analyzed using two-way ANOVA followed by Tukey’s multiple-comparisons test; exact P values are indicated. Cytokine concentrations in **(G,H)** were log₂-transformed and normalized to the geometric mean of the corresponding LPS group. Data in **(H)** are presented as mean ± SD with individual values. Statistical differences were assessed using the Kruskal–Wallis test followed by Dunn’s multiple-comparisons test against LPS control. * indicates adjusted P < 0.05.

To characterize tissue-level responses in the corresponding organs, H&E-stained sections were examined (**Figure 6D**). Liver sections from pITA-NP- and pMSA-NP-treated mice showed broadly preserved hepatic architecture, with recognizable hepatocyte plates, sinusoidal spaces, and vascular structures and no conspicuous confluent necrosis or extensive inflammatory aggregates in the examined fields. The LPS-only liver showed mild hepatocellular cytoplasmic clearing and focal vascular/sinusoidal congestion, although overall hepatic architecture remained recognizable. As shown in **Figure 6D**, the LPS lung showed increased alveolar septal and parenchymal cellularity; the kidney showed mild tubular epithelial alterations with focal vascular congestion; and the spleen showed greater heterogeneity in white-pulp organization, whereas myocardial architecture remained broadly preserved. Corresponding sections from the nanoparticle-treated groups retained readily identifiable tissue architecture across these organs.

To complement the histological observations, circulating ALT, AST, BUN, and creatinine were measured to assess hepatic and renal biochemical responses during LPS-induced systemic inflammation (**Figure S13**). No consistent nanoparticle-associated elevations were observed across these markers. Notably, pMSA-NP treatment was associated with numerically lower ALT and AST levels relative to the LPS control, whereas BUN and creatinine did not exhibit a consistent treatment-associated reduction. Together, the histological and biochemical findings are consistent with short-term organ tolerability of both nanoparticle formulations under the tested LPS-challenge conditions.

Nanoparticle treatment was associated with higher survival following LPS challenge (**Figure 6E,F**), with seven-day survival rates of 15.4 % for vehicle (n = 13), 50 % for pITA-NP (n = 10), and 40 % for pMSA-NP (n = 10). However, neither nanoparticle treatment reached statistical significance after adjustment for multiple comparisons (adjusted P = 0.1204 and 0.1793, respectively; log-rank test). He et al. previously reported significantly prolonged survival with free mesaconate (250 mg kg^-1^, 2 h before a 30 mg kg^-1^ LPS challenge over 24 h).^10^ In contrast, the present study used a degradable pMSA nanoparticle 3 h before a 20 mg kg^-1^ LPS challenge with seven-day monitoring. Although the survival difference observed with pMSA-NP did not reach statistical significance and the different LPS doses, treatment regimens, sample sizes, and observation periods preclude direct efficacy comparison, the present platform extends MSA from a soluble immunometabolite into an engineerable nanomaterial capable of systemic cytokine modulation and preliminary survival-associated activity *in vivo*.

Circulating cytokines and chemokines were further evaluated using a 26-plex Luminex assay, with normalized responses shown in **Figure 6G,H** and corresponding concentrations in **Figures S14** and **S15**. Both nanoparticle formulations reduced multiple LPS-responsive inflammatory mediators. The largest response was observed for IL-6, with mean log₂ fold changes of −1.27 and −3.38 for pITA-NP and pMSA-NP, respectively, corresponding to approximately 2.4- and 10.4-fold lower geometric mean concentrations relative to LPS control. IL-6 was significantly reduced by both formulations, with pMSA-NP also lower than pITA-NP (**Figure S14D**), consistent with modulation of the IL-6/JAK/STAT3-associated signaling identified transcriptionally (**Figure 5G**).^52^

Additional inflammatory mediators exhibited overlapping but quantitatively different responses to the two nanoparticle formulations. pMSA-NP showed greater reductions in TNF-α, MCP-1/CCL2, and IL-12p70 relative to LPS control, with similarly strong decreases observed for several additional inflammatory mediators, including IL-17A, IL-18, IL-23, GRO-α/CXCL1, and MIP-family chemokines (**Figure S14**). In contrast, pITA-NP produced a greater reduction in IP-10/CXCL10 (−0.90 versus −0.59 log_2_ fold change for pITA-NP and pMSA-NP, respectively), with the pITA-NP-associated reduction reaching statistical significance relative to LPS (**Figure 6H**). Given the interferon-associated transcriptional changes identified in macrophages, this finding provides a complementary systemic readout of inflammatory signaling. CXCL10 is an interferon-inducible CXCR3 ligand that is closely linked to type I and type II interferon responses and inflammatory leukocyte trafficking.^53,54^ Interestingly, He et al. reported increased CXCL10 following soluble MSA treatment, whereas pMSA-NP pretreatment was associated with lower circulating IP-10/CXCL10 relative to LPS control.^10^ Although these findings arise from different experimental contexts, they suggest that polymer-based MSA likely produces a biological profile distinct from the free metabolite. Together, these results suggest that pITA-NP and pMSA-NP elicit overlapping but potentially distinguishable systemic cytokine responses, although direct differences between the formulations were not statistically established.

The circulating response also extended beyond conventional inflammatory mediators. IL-10 decreased following pITA-NP and pMSA-NP pretreatment, together with reductions across several type 2-associated and context-dependent cytokines, including IL-4, IL-5, IL-9, IL-13, IL-22, and IL-27 (**Figure S15**), whereas IL-2 showed no clear treatment-associated separation. These changes indicate broader remodeling of the LPS-induced cytokine milieu rather than selective enhancement of anti-inflammatory cytokine production. IL-10 is a well-established immunoregulatory cytokine, whereas IL-4 and IL-13 are canonical type 2 cytokines with context-dependent roles in macrophage activation, tissue repair, and inflammation.^55,56^ Together with the macrophage phenotypic and transcriptomic findings, the systemic cytokine profiles are consistent with non-canonical immunomodulation involving inflammatory, regulatory, and interferon-associated signaling. Because the nanoparticles were administered before LPS challenge, the survival findings should be interpreted within a prophylactic treatment paradigm rather than as evidence of post-challenge therapeutic efficacy. Although the treatment-associated differences in 7-day survival did not reach statistical significance after multiple-comparison adjustment, both nanoparticle formulations were associated with numerically higher survival relative to vehicle control. This finding is consistent with emerging evidence that polymerized itaconate-based nanoparticles can influence outcomes during severe systemic inflammatory challenge; a recent pITA nanoparticle study similarly reported increased survival in experimental sepsis together with reduced inflammation and multiorgan injury.^57^ In the present study, the distinct material organization, macrophage transcriptional programs, systemic cytokine-response profiles, and survival outcomes collectively suggest that metabolite-isomer selection may diversify the biological responses accessible through intrinsically bioactive polyester nanoparticles.

### 2.7. Materials-to-Biology Framework for Metabolite-Based Nanoparticle Design

Integration of the materials, macrophage, transcriptomic, and *in vivo* findings supports a structure-process-property-bioactivity framework (**Scheme 1**) in which metabolite-isomer geometry and synthesis conditions generate distinct nanoparticle material states that may influence degradation-associated accessibility and immunometabolic responses. Beyond the previously established immunomodulatory activity of soluble MSA,^10^ the present study establishes pMSA as a distinct, synthesis-tunable building block for immunometabolically-active biomaterials. Incorporation of MSA into a degradable polyester introduces material-level design variables, including molecular-weight development, thermal organization, nanoparticle formulation, and degradation-associated accessibility, that are absent from the free metabolite. Building on prior advances in pITA-based polyester systems,^21,32^ pMSA therefore emerges as a distinct, synthesis-tunable building block for immunometabolically active biomaterials and expands the material and biological response space available for cargo-free immunometabolic design.

Collectively, these findings position mesaconate-based polyester nanoparticles as a synthesis-tunable, intrinsically bioactive platform for cellular and systemic immunomodulation. More broadly, it demonstrates that endogenous immunometabolites can function not only as soluble signaling molecules but also as structural components of degradable biomaterials, enabling metabolite identity, polymer organization, and biological accessibility to be engineered together. This materials-based framework broadens the design space of immunometabolic therapeutics beyond conventional soluble analogs and provides a foundation for developing metabolite-based nanomaterials for inflammatory disease modulation.

**Scheme 1.**
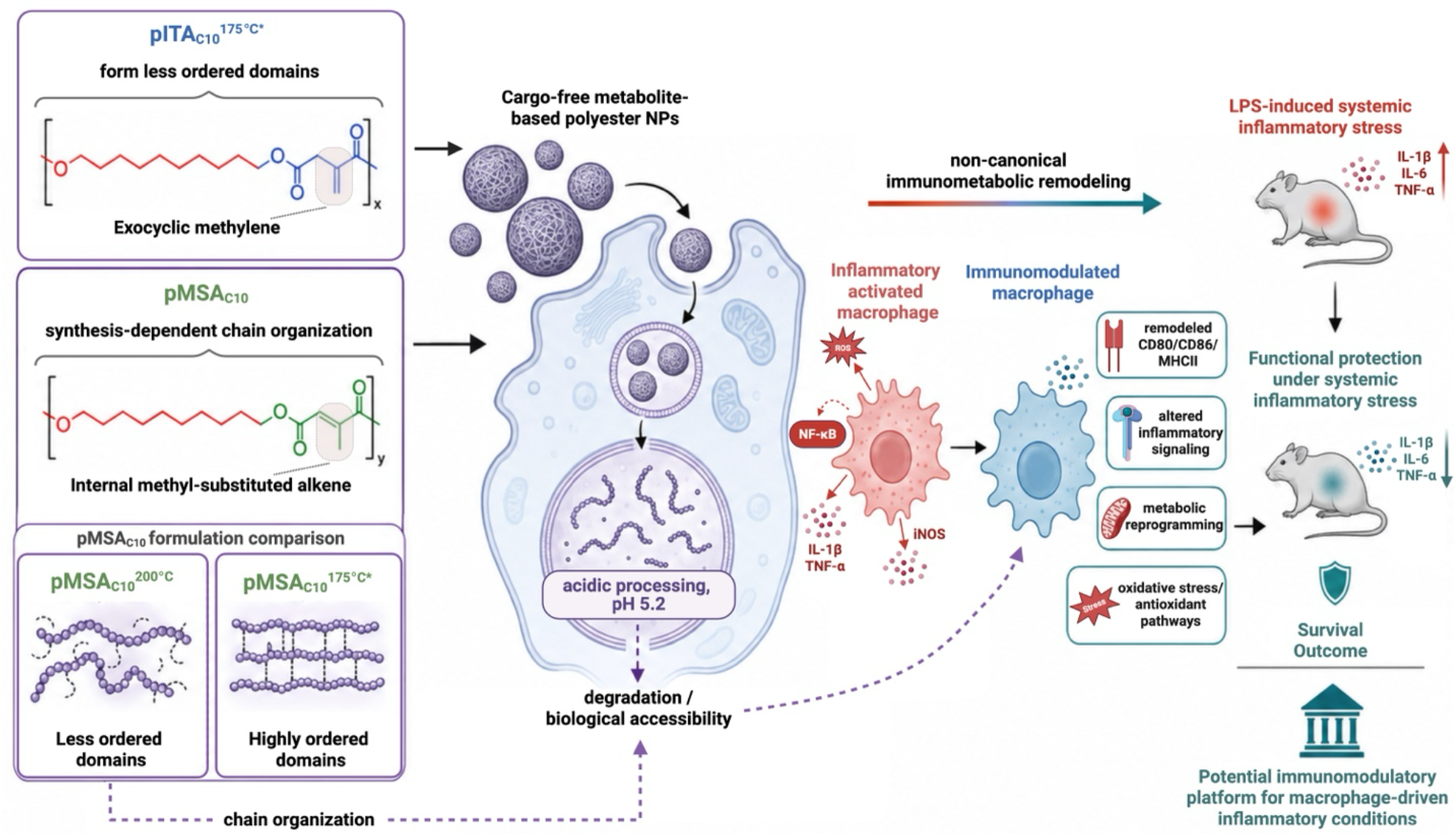
Proposed materials-to-biology framework for metabolite-based polyester nanoparticles

Metabolite-isomer structure and synthesis-controlled polymer organization influence nanoparticle properties and degradation-associated accessibility, potentially shaping macrophage immunometabolic remodeling. Associated systemic cytokine modulation and improved endotoxemia survival support the potential of cargo-free metabolite-derived nanoparticles for inflammatory conditions. Dashed arrows indicate proposed mechanistic links.

## 3. Conclusion

This study establishes MSA-based polyester nanoparticles as a synthesis-tunable platform for macrophage-directed immunomodulation. By comparing structurally related ITA- and MSA-derived polyesters, we show that metabolite isomerism and synthesis conditions influence molecular-weight development, thermal organization, degradation-associated behavior, and nanoparticle bioactivity. Although pITA-NP and pMSA-NP elicited substantially overlapping transcriptional responses in unstimulated macrophages, their responses diverged markedly under LPS stimulation. pITA-NP preferentially engaged redox-, metabolic-, and cellular-stress-associated programs, whereas pMSA-NP produced a more distinct inflammatory-, remodeling-,and lysosome-associated transcriptional profile, demonstrating that the two metabolite-derived materials access different macrophage response states despite their structural similarity. Both formulations also produced systemic cytokine modulation and were associated with higher 7-day survival following LPS challenge relative to vehicle control. Using MSA as a monomer to construct a degradable polyester transforms an underexplored soluble immunometabolite into an engineerable material in which metabolite identity, polymer organization, degradation-associated accessibility, and biological response can be tuned together. More broadly, these findings position mesaconate as a distinct building block for intrinsically bioactive biomaterials and expand the design space of cargo-free immunometabolic interventions beyond established ITA-based systems.

## 4. Experimental Section

### 4.1 Materials

Itaconic acid, mesaconic acid, 1,10-decanediol, 4-octyl itaconate (4OI), 4-methoxyphenol (MeHQ), tin(II) chloride (SnCl₂), poly(vinyl alcohol) (PVA; 87–90% hydrolyzed; average molecular weight 30,000–70,000), chloroform, sucrose, d-mannitol, LPS (*Escherichia coli* O111:B4, #L2630), and sodium acetate buffer were purchased from MilliporeSigma (Burlington, MA, USA). RPMI 1640 supplemented with L-glutamine and penicillin/streptomycin was purchased from Invitrogen (Waltham, MA, USA), and heat-inactivated fetal bovine serum (FBS) was purchased from VWR (Radnor, PA, USA). L929 cells were obtained from ATCC (Manassas, VA, USA). FITC Annexin V Apoptosis Detection Kit with PI was purchased from BioLegend (San Deigo, CA, USA). Additional antibodies and biological reagents are identified in the corresponding experimental sections.

### 4.2 Synthesis and purification of pITA_C10_ and pMSA_C10_ polyesters

Poly(itaconate-co-decanediol) (pITA_C10_) and poly(mesaconate-co-decanediol) (pMSA_C10_) were synthesized by catalyst-assisted melt polycondensation of the corresponding dicarboxylic acid with 1,10-decanediol. Briefly, itaconic acid or mesaconic acid (650.05 mg, 5.0 mmol, 1.0 equiv.) and 1,10-decanediol (871.5 mg, 5.0 mmol, 1.0 equiv.) were added to a dry 25 mL round-bottom flask equipped with a magnetic stir bar. For inhibitor-containing formulations, MeHQ (7 mg, 0.056 mmol, 0.011 equiv.) was added before catalyst addition. The flask was sealed, purged with argon, and heated to the designated reaction temperature until the reaction mixture was fully molten and homogeneous. SnCl₂ (107 mg, 0.56 mmol, 0.11 equiv.) was then added, and polymerization was performed for 2 h, consisting of 1 h under argon followed by 1 h under in-house vacuum.

Four polymer candidates were prepared: pITA_C10_^175^ ^°C*^, pMSA_C10_^200^ ^°C^, pMSA_C10_^175^ ^°C^, and pMSA_C10_^175^ ^°C*^. where the asterisk denotes synthesis in the presence of MeHQ. At the reaction endpoint, the flask was immediately quenched in an ice bath. The crude polymer was dissolved in CHCl₃ (8 mL) and precipitated dropwise into cold, stirred methanol (300 mL). The precipitated polymer was collected by vacuum filtration and dried under vacuum for 24–48 h before characterization and nanoparticle formulation. Formulation-specific conditions are summarized in **Table S1**. For polymerization-kinetics experiments, molecular-weight development was evaluated at 0.5, 1.0, 1.5, 2.0, and 2.5 h. Reactions were conducted using the same formulation-specific synthesis conditions described above, with the first 1 h under argon followed by house vacuum for the remainder of the reaction. Aliquots were withdrawn from the ongoing reaction at each designated time point, quenched immediately in ice-bath, and subsequently analyzed by GPC. Each formulation was evaluated using three independent synthesis batches (n = 3).

### 4.3 Nanoparticle formulation

Nanoparticles were formulated using an oil-in-water single-emulsion solvent evaporation method. Briefly, each polymer was dissolved in chloroform at 50 mg mL^-1^ using a 100 mg polymer batch and allowed to dissolve overnight. Any undissolved material was removed by centrifugation (12,000× g at 4 °C for 10 min), and the clarified polymer solution was transferred to a 20 mL scintillation vial. An aqueous 1 % poly(vinyl alcohol) (PVA) solution, 10 mL, was slowly added to the polymer solution to form a two-phase system. The mixture was emulsified on ice using a Cole-Parmer 500 W ultrasonic homogenizer equipped with a 1/8^th^ inch probe tip for 30 s at 40 % amplitude. Immediately after sonication, the emulsion was poured into 80 mL of stirring 0.5 % PVA and stirred overnight to allow chloroform evaporation and nanoparticle hardening. The resulting nanoparticle suspension was filtered through a 40 µm cell strainer to remove large aggregates. Nanoparticles were then washed three times with 40 mL cold deionized water by centrifugation at 12,000× g for 20 min. The final nanoparticle pellet was resuspended in cryoprotectant solution containing 4 % w/v sucrose and 3 % w/v mannitol, aliquoted into 2 mL screw-cap microcentrifuge tubes, frozen at −80 °C, and lyophilized for 2 days. Similarly, Cy5.5-labeled NPs were formulated using synthesized Cy5.5-polymer conjugates as previously reported.^58^ Cy5.5-polymer conjugate was incorporated using 0.5 % (w/w) conjugate into the NP formulations.

### 4.4 Polymer and nanoparticle physicochemical characterization

Polymer molecular weight and dispersity were determined by gel permeation chromatography (GPC). Dried polymer samples were dissolved in chloroform at 5 mg mL^-1^ overnight, filtered through a 0.2 µm syringe filter, and analyzed using a Waters Acquity HPLC system equipped with Styragel HR4E and HR3 columns connected in series. Chloroform was used as the mobile phase at a flow rate of 1 mL min^-1^. Molecular-weight measurements were performed using triple detection with a Malvern Panalytical OMNISEC detector, and M_n_, M_w_, and dispersity (Đ = M_w_/M_n_) were calculated using OMNISEC v11.41 software.

Polymer chemical structures were characterized by ^1^H NMR and ATR-FTIR spectroscopy. For ^1^H NMR analysis, dried polymers were dissolved in CDCl_3_ at 15 mg mL^-1^, filtered through a 2.0 µm syringe filter, and analyzed using a Varian 400-MR spectrometer at 25 °C. Spectra were acquired using the s2pul pulse sequence with 8 scans and a 5 s relaxation delay. Chemical shifts were referenced to residual CHCl_3_ at δ 7.26 ppm. ATR-FTIR spectra were collected using a JASCO FTIR-6100 spectrometer equipped with a diamond ATR accessory. Approximately 5 mg of dried polymer was placed directly on the ATR crystal, and spectra were acquired in absorbance mode over 4000–400 cm^-1^.

Thermal properties of bulk polymers and lyophilized nanoparticles were analyzed by differential scanning calorimetry using a DSC2500 instrument. Approximately 4 mg of sample was placed in a standard aluminum DSC pan and analyzed under an argon purge. Samples were equilibrated at −70 °C, heated to 70 °C at 10 °C min^-1^, held isothermally for 2 min, cooled to −70 °C at 10 °C min^-1^, held isothermally for 2 min, and reheated to 70 °C at 10 °C min^-1^. Reported thermal transitions were determined from the second heating cycle.

Nanoparticle hydrodynamic diameter, polydispersity index (PDI), and zeta potential were measured using a Zetasizer Nano ZSP (Malvern Panalytical, Worcestershire, UK). Hydrodynamic diameter and PDI were determined by dynamic light scattering, whereas zeta potential was determined from electrophoretic mobility. Samples were diluted 1:100 in dH_2_O and measured in triplicate at 25 °C.

Nanoparticle morphology was examined by scanning electron microscopy. Nanoparticle suspensions were prepared at 1 mg mL^-1^, and 2 µL of each suspension was deposited onto an SEM stub and dried. Samples were gold sputter-coated before imaging. SEM images were acquired using an Apreo 2S VS instrument operated at 2.5 kV and a beam current of 0.10 nA, with magnifications ranging from 30,000× to 80,000×.

### 4.5 Acidic incubation and degradation-associated nanoparticle stability

To evaluate formulation-dependent behavior under acidic conditions relevant to endosomal/lysosomal processing, lyophilized pITA_C10_^175^ ^°C*^-NPs, pMSA_C10_^200^ ^°C^-NPs, pMSA_C10_^175^ ^°C^-NPs, and pMSA_C10_^175^ ^°C*^-NPs were dispersed in pH 5.2 acetate buffer at a nanoparticle concentration of 3 mg mL^-1^ and incubated at 37 °C under gentle agitation. Samples were evaluated at 0 h, 24 h, 7 d, and 14 d. Hydrodynamic diameter and PDI were measured by DLS at each time point.

For GPC analysis, nanoparticles were recovered by centrifugation (12,000× g at 4 °C for 10 min), washed twice with deionized water to remove buffer salts, frozen, lyophilized, dissolved in chloroform, filtered, and analyzed using the GPC method described above. M_n_ retention was calculated relative to the formulation-specific recovered-polymer baseline at 0 h according to:

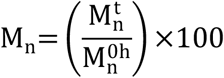

These measurements were used to assess degradation-associated molecular-weight changes and colloidal restructuring rather than complete polymer degradation.

### 4.6 Bone marrow-derived macrophages (BMMØs) isolation, culture, and NP treatment

C57BL/6J male mice, 5–7 weeks of age, were obtained from The Jackson Laboratory (Bar Harbor, ME, USA) and maintained under pathogen-free conditions at the University of Maryland, Baltimore. All animal procedures were approved by the University of Maryland, Baltimore Institutional Animal Care and Use Committee (IACUC protocol no. AUP-00000975). Bone marrow was isolated from femurs and tibias and flushed with macrophage differentiation medium comprising RPMI 1640 supplemented with L-glutamine, penicillin (100 units mL^-1^), streptomycin (100 µg mL^-1^), 10 % FBS, and 20 % L929 cell-conditioned medium. Red blood cells were removed using ACK lysis buffer, and the resulting suspension was passed through a 40 µm cell strainer. Cells were counted by trypan-blue exclusion, plated at 2 × 10^6^ cells per 10 cm tissue-culture-treated dish, and maintained at 37 °C and 5 % CO_2_. Medium was replaced on days 3, 6, and 8, and differentiated macrophages were used on days 7–10.

### 4.7 Nanoparticle uptake

Macrophages were seeded at 30,000 cells per well in an 8-well Millicell EZ glass chamber slide (MilliporeSigma, Burlington, MA, USA). Cells were treated with 30 µg mL^-1^ Cy5.5-labeled nanoparticles for 24 h. After treatment, cells were washed with 1x HBSS, fixed with fixation buffer, and stained with wheat germ agglutinin (WGA)-Alexa Fluor 594 conjugate (Thermo Fisher Scientific, Waltham, MA, USA). After staining, cells were mounted using Fluoroshield with DAPI (Sigma-Aldrich, St. Louis, MO, USA). Images were taken at 20× magnification using a Nikon Eclipse TI using NIS-Elements 5.42.06 software (Nikon, Melville, NY, USA).

### 4.8 Macrophage viability and flow-cytometric phenotyping

Bone marrow-derived macrophages were seeded at 0.2 × 10^6^ cells per well in 24-well plates. Where indicated, inflammatory activation was established by overnight stimulation with LPS (100 ng mL^-1^). Cells were subsequently treated for 24 h with nanoparticles (30 µg mL^-1^) or 4OI (125 µM).

For viability and surface-marker phenotyping, cells were stained with Live/Dead Green Viability/Cytotoxicity dye (Thermo Fisher Scientific, Waltham, MA, USA), followed by Fc-receptor blocking with anti-CD16/32 (clone 93). Cells were then stained with antibodies against F4/80 (clone BM8, PE/Cyanine7), CD11b (clone M1/70, Pacific Blue), CD80 (clone 16-10A1, Brilliant Violet 510), CD86 (clone GL-1, APC/Cyanine7), and MHCII/I-A/I-E (clone M5/114.15.2, PE-Fire 640), all purchased from BioLegend (San Diego, CA, USA). Samples were fixed and acquired using a Cytek Aurora 3 flow cytometer and analyzed using FCS Express 7 (De Novo Software, Glendale, CA, USA). Macrophages were identified based on CD11b and F4/80 expression, and CD80, CD86, and MHCII expression was quantified as median fluorescence intensity as shown by the gating strategy demonstrated in **Figure S6**. For apoptosis analysis, cells were stained with F4/80, CD11b, FITC-Annexin V, and propidium iodide and analyzed immediately without fixation. Annexin V^-^/PI^-^ cells were classified as live, Annexin V^+^/PI^-^ cells as early apoptotic, and Annexin V^+^/PI^+^ cells as dead/late-stage apoptotic.

### 4.9 RNA sequencing and bioinformatic analysis

RNA isolation, sequencing, and primary data processing were performed by the University of Maryland Institute for Genome Sciences (IGS). BMDMs maintained under resting conditions or stimulated with LPS were treated with pITA-NPs or pMSA-NPs (30 µg mL^-1^) for 24 h; soluble 4OI (125 µM) was included as a comparator. IL-4 stimulated macrophages served as an alternatively activated macrophage control. Gene-level count matrices and DESeq2 differential-expression outputs generated by IGS were used for downstream analyses. Treatment groups were compared with the corresponding untreated M0 or LPS control, and differentially expressed genes were defined by |log_2_ fold change| ≥ 0.58 and Benjamini–Hochberg-adjusted p < 0.05. Downstream analyses were performed in R v4.6.1 and included principal component analysis, volcano plots, Pearson correlation of pITA-NP and pMSA-NP transcriptional responses, targeted heatmaps, and gene set enrichment analysis using KEGG and MSigDB Hallmark gene sets. Enrichment results were summarized by normalized enrichment score and Benjamini–Hochberg-adjusted false discovery rate.

### 4.10 LPS-induced endotoxemia survival study

Male C57BL/6J mice (6-8 weeks) were randomly assigned to treatment groups and administered pITA-NP or bioactivity-selected pMSA-NP by intraperitoneal (i.p.) injection (2 mg per mouse; 10 mg mL⁻¹; n = 10 per nanoparticle group) or 200 µL saline vehicle (n = 13) 3 h before i.p. LPS challenge (20 mg kg⁻¹). Animals were monitored for survival for 7 days.

### 4.11 In vivo biodistribution and inflammation study

A separate cohort of male C57BL/6J mice was used to evaluate the effects of pITA-NP and bioactivity-selected pMSA-NP during LPS-induced inflammation. Mice received 2 mg kg^-1^ Cy5.5-NPs (5 mice per nanoparticle group) or 200 µL saline for control groups (3 mice per NT and LPS group) i.p. 3 h prior to an i.p. LPS challenge (5 mg kg^-1^). After 3 h of LPS treatment, the blood plasma, peritoneal fluid, and organs were collected for downstream analysis.

#### Biodistribution

Nanoparticle-associated fluorescence was quantified ex vivo in the heart, lungs, spleen, liver, and kidneys, with representative organ images and quantitative fluorescence measurements used to evaluate organ-level distribution. Organs were analyzed on the Xnogen in vivo imaging system (IVIS) Spectrum Optical (PerkinElmer, Waltham, MA, USA) with excitation and emission wavelengths of 675 and 720 nm respectively.

#### Blood chemistry

Circulating ALT, AST, BUN, and creatinine were quantified to assess short-term hepatic and renal biochemical responses. Blood plasma was sent for analysis by VRL Diagnostics (Gaithersburg, MD, USA).

#### Circulating cytokines and chemokines

Circulating mediators were quantified using a 26-plex Luminex panel (EPXR260-26088-901) and processed using the Luminex xPONENT software (Thermo Fisher Scientific, Waltham, MA, USA). Cytokine concentrations were analyzed both as measured concentrations and after log_2_ normalization relative to the geometric mean of the corresponding LPS control group for each analyte.

#### H&E staining

Organ damage and alterations in tissue-level architecture was assessed via histopathological evaluation. Following IVIS, organs were rinsed in PBS, placed into tissue cassettes, and fixed for 24 h in 10 % buffered formalin (Sigma-Aldrich, St. Louis, MO). The following day, samples were rinsed with PBS and delivered to the Pathology Biorepository Shared Services Core at the University of Maryland, Baltimore, where the samples were paraffin embedded and sectioned for hematoxylin and eosin (H&E) staining.

### 4.12 Statistical Analysis

Statistical analyses were performed using GraphPad Prism v11.1.0 (GraphPad Software, Boston, MA, USA), unless otherwise specified. Data are presented as mean ± SD, with individual data points shown where applicable. For cell-based experiments in Figure 4, comparisons among multiple treatment groups were performed using one-way ANOVA followed by Dunnett’s multiple-comparisons test against the corresponding control, whereas selected pairwise comparisons were analyzed using two-sided unpaired t-tests with Welch’s correction. For biodistribution analyses, two-way ANOVA followed by Tukey’s multiple-comparisons test was used. Blood-chemistry markers, including ALT, AST, BUN, and creatinine, were analyzed using one-way ANOVA followed by Dunnett’s multiple-comparisons test against the corresponding control group. Circulating cytokine and chemokine data were analyzed using the Kruskal–Wallis test followed by Dunn’s multiple-comparisons test for the indicated comparisons. Survival distributions were compared using the log-rank (Mantel–Cox) test, with each nanoparticle-treated group compared with the vehicle control and Holm–Šídák correction applied for multiple comparisons. Multiplicity-adjusted P values were reported using a familywise α of 0.05. For RNA-seq analyses, multiple-testing correction was performed using the Benjamini–Hochberg procedure as described in Section 4.9. Statistical significance was defined as P < 0.05 or adjusted P < 0.05, as appropriate. Exact P values are reported where applicable.

The definition of n was experiment-specific and is provided in the corresponding figure or table legend. For polymer-synthesis and polymerization-kinetics studies, n represents independently prepared synthesis batches. For acidic-incubation GPC and DLS analyses, n = 3 measurements per formulation and time point. For cell-based studies, n represents independent biological experiments; RNA-seq was performed using n = 3 independent biological replicates per condition; and for *in vivo* studies, n represents individual animals.

## Supporting Information

Supporting Information is available from…

## Acknowledgements

This work was supported by the National Institute of General Medical Sciences (R35GM142752) and the National Institute of Allergy and Infectious Diseases (R01AI192405), awarded to R.M.P., and by the National Institute of Diabetes and Digestive and Kidney Diseases (R01DK144500), awarded to A.H.N., all of the National Institutes of Health. This publication was supported by funds through the Maryland Department of Health’s Cigarette Restitution Fund Program and the National Cancer Institute - Cancer Center Support Grant (CCSG) - P30CA134274; University of Maryland School of Medicine’s Center for Translational Research in Imaging - Baltimore, Maryland; University of Maryland School of Medicine Genomics Core - Baltimore, Maryland; University of Maryland Pathology Biorepository Shared Services - Baltimore, Maryland; University of Maryland School of Medicine’s & Greenebaum Comprehensive Cancer Center’s Flow Cytometry Core – Baltimore, Maryland; and University of Maryland School of Medicine’s and School of Dentistry’s Electron Microscopy Core - Baltimore, Maryland. The content is solely the responsibility of the authors and does not necessarily represent the official view of the National Institutes of Health.

## Data Availability Statement

The data that support the findings of this student are available from the corresponding author upon reasonable request.

