## Supplementary Information for "Mesaconate-Based Polyester Nanoparticles Reveal How Metabolite Isomerism Shapes Polymer Behavior, Macrophage Reprogramming, and Systemic Immunomodulation"

### 1. Materials

Itaconic acid, mesaconic acid, 1,10-decanediol, 4-octyl itaconate (4OI), 4-methoxyphenol (MeHQ), tin(II) chloride ($SnCl_2$), poly(vinyl alcohol) (PVA; 87–90 % hydrolyzed; average molecular weight 30,000–70,000), chloroform, sucrose, d-mannitol, LPS (*Escherichia coli* O111:B4, #L2630) and sodium acetate buffer were purchased from MilliporeSigma (Burlington, MA, USA). Acid-terminated poly(D,L-lactide-co-glycolide) (PLGA), of low inherent viscosity in hexafuoro-2-propanonal ~ 0.17 dL/g (approximate MW 4200 g/mol) was purchased from Lactel® Absorbable Polymers (Birmingham, AL., USA). RPMI 1640 supplemented with L-glutamine and penicillin/streptomycin was purchased from Invitrogen (Waltham, MA, USA). Heat-inactivated fetal bovine serum (FBS) was purchased from VWR (Radnor, PA, USA), and L929 cells were obtained from ATCC (Manassas, VA, USA). FITC Annexin V Apoptosis Detection Kit with PI was purchased from BioLegend (San Deigo, CA, USA).

### 2. Synthesis and purification of $pITA_{C10}$ and $pMSA_{C10}$ polyesters

Poly(itaconate-co-decanediol) ($pITA_{C10}$) and poly(mesaconate-co-decanediol) ($pMSA_{C10}$) were synthesized by catalyst-assisted melt polycondensation of the corresponding unsaturated dicarboxylic acid with 1,10-decanediol. The same reaction scale and general procedure were used for all polymer candidates, while acid-monomer identity, reaction temperature, and MeHQ inhibitor use were varied as summarized in **Table S1**.

A dry 25 mL round-bottom flask equipped with a magnetic stir bar was charged with itaconic acid or mesaconic acid (650.05 mg, 5.0 mmol, 1.0 equiv.) and 1,10-decanediol (871.5 mg, 5.0 mmol, 1.0 equiv.). For inhibitor-containing formulations, MeHQ (7 mg, 0.056 mmol, 0.011 equiv.) was added before catalyst addition. The flask was sealed with a rubber septum and purged with argon using a needle-in/needle-out configuration to remove air from the reaction vessel. The flask was placed in a preheated heating bath at the designated reaction temperature and maintained under argon until the monomer, diol, and inhibitor, where present, became fully molten and homogeneous. $SnCl_2$ (107 mg, 0.56 mmol, 0.11 equiv.) was then added as the polycondensation catalyst. The flask was immediately resealed and purged again with argon, and the reaction timer was initiated. Polymerization was performed for 2 h, consisting of 1 h under argon followed by 1 h under laboratory house vacuum. The vacuum pressure was not independently measured.

Four polymer formulations were prepared: $pITA_{C10}^{175\ °C*}$, $pMSA_{C10}^{200\ °C}$, $pMSA_{C10}^{175\ °C}$, and $pMSA_{C10}^{175\ °C*}$, where the asterisk denotes synthesis in the presence of MeHQ. The corresponding formulation-specific conditions are summarized in **Table S1**.

At the reaction endpoint, the flask was removed from the heating bath and immediately quenched in an ice bath to stop polymerization. The crude polymer was dissolved in $CHCl_3$ (8 mL) to obtain a homogeneous polymer solution. The solution was added dropwise into cold, stirred methanol (300 mL) to precipitate the polymer and remove unreacted monomers and low-molecular-weight oligomers. The precipitation mixture was stirred until the supernatant became clear. The precipitated polymer was collected by vacuum filtration using a Büchner funnel and filter paper, transferred to a pre-weighed scintillation vial, and dried under vacuum for 24–48 h.

The dried polymer was weighed to determine isolated yield and stored until physicochemical characterization and nanoparticle formulation.

#### *2.1 Polymerization-kinetics analysis*

Polymer molecular-weight development was evaluated at reaction times of 0.5, 1.0, 1.5, 2.0, and 2.5 h. Reactions were conducted using the same formulation-specific synthesis conditions described above. The first 1 h was performed under argon, followed by house vacuum for the remainder of the reaction. Aliquots were withdrawn from the ongoing reaction at the designated time points, immediately quenched in an ice bath to arrest further polymerization, and subsequently prepared for GPC analysis. Each formulation was evaluated using three independent synthesis batches (n = 3).

### 3. Nanoparticle formulation

Nanoparticles were prepared using an oil-in-water single-emulsion solvent-evaporation method. Each polymer (100 mg) was dissolved in chloroform at 50 mg mL-1 and allowed to dissolve overnight. Any undissolved material was removed by a 10 min, 12,000 x g centrifugation at 4 °C. The clarified polymer solution was then transferred to a 20 mL scintillation vial. An aqueous 1 % (w/v) PVA solution (10 mL) was slowly added to the polymer solution to form a two-phase system. The mixture was emulsified on ice for 30 s at 40 % amplitude using a Cole-Parmer 500 W ultrasonic homogenizer equipped with a 1/8$^{th}$-inch probe tip.

Immediately after sonication, the emulsion was transferred to 80 mL of stirring 0.5 % (w/v) PVA and stirred overnight to allow chloroform evaporation and nanoparticle hardening. The resulting nanoparticle suspension was passed through a 40 µm cell strainer to remove large aggregates. Nanoparticles were washed three times with 40 mL cold deionized water by centrifugation at 12,000 × g for 20 min. The final nanoparticle pellet was resuspended in cryoprotectant solution containing 4 % (w/v) sucrose and 3 % (w/v) mannitol, aliquoted into 2 mL screw-cap microcentrifuge tubes, frozen at −80 °C, and lyophilized for 2 days. Similarly, Cy5.5-labeled NPs were formulated using synthesized Cy5.5-polymer conjugates as previously reported. In brief, PLGA was first conjugated to cyanine 5.5 (Lumiprobe, Cockeysville, MD, USA) using EDC/NHS carbodiimide chemistry. The resulting product was then transferred to a 3.5 kD dialysis bag and dialyzed against 4 L of $dH_2O$ for 48 h, with 6 water exchanges. The purified conjugate was then lyophilized for 2 days. The resulting Cy5.5-polymer conjugate was incorporated using 0.5 % (w/w) conjugate into the NP formulations.

### 4. Physicochemical characterization

#### *4.1 Gel permeation chromatography*

The molecular weight and dispersity of synthesized polyesters were determined by gel permeation chromatography (GPC). Dried polymer samples were dissolved in chloroform at 5 mg mL$^{-1}$ and allowed to dissolve overnight at room temperature. Before injection, samples were filtered through a 0.2 μm syringe filter to remove undissolved particulates.

GPC analysis was performed using a Waters Acquity HPLC system equipped with Styragel HR4E and HR3 columns connected in series and maintained at 25 °C. Chloroform was used as the mobile phase at a flow rate of 1 mL min$^{-1}$. Samples were analyzed using injection volumes of 75–100 μL. Molecular-weight measurements were performed using triple detection with a Malvern Panalytical OMNISEC detector. Number-average molecular weight ($M_n$), weight-average molecular weight ($M_w$), and dispersity ($Đ = M_w/M_n$) were calculated using Malvern OMNISEC v11.41 software.

#### *4.2 $^1$H NMR spectroscopy*

$^1$H NMR spectroscopy was used to confirm polyester formation and retention of characteristic itaconate- or mesaconate-derived backbone proton resonances. Dried polymer samples were dissolved in $CDCl_3$ at 15 mg mL$^{-1}$ and filtered through a 2.0 μm syringe filter before analysis. Spectra were acquired using a Varian 400-MR spectrometer at 25 °C using the s2pul pulse sequence with 8 scans and a 5 s relaxation delay. Chemical shifts were reported in ppm relative to residual $CHCl_3$ at δ 7.26 ppm.

The spectra were used to identify characteristic proton resonances corresponding to the C10 aliphatic linker and ITA- or MSA-derived unsaturated backbone units. Spectral assignments were used to confirm structure rather than to quantitatively determine alkene retention.

#### *4.3 ATR-IR spectroscopy*

Attenuated total reflectance infrared spectroscopy was performed to evaluate polyester formation and backbone-associated chemical features. Dried polymer samples, approximately 5 mg, were placed directly onto the diamond ATR crystal of a JASCO FTIR-6100 spectrometer equipped with a diamond ATR accessory. Spectra were collected in absorbance mode. Spectra were acquired over the range of 4000 – 400 cm$^{-1}$, with a data interval of 0.964 cm$^{-1}$ and 3736 data points per spectrum. The resulting spectra were used to identify characteristic ester carbonyl stretching, ester C–O–C stretching, aliphatic C–H stretching, and alkene/backbone-associated vibrational features.

#### *4.4 Differential scanning calorimetry*

Thermal properties of the bulk polymers and lyophilized nanoparticles were analyzed using a DSC2500 differential scanning calorimeter. Approximately 4 mg of each sample was placed in a standard aluminum DSC pan and analyzed under an argon purge.

Samples were equilibrated at −70 °C, heated to 70 °C at 10 °C min$^{-1}$, held isothermally for 2 min, cooled to −70 °C at 10 °C min$^{-1}$, held isothermally for 2 min, and reheated to 70 °C at 10 °C min$^{-1}$. Reported glass-transition, melting, and cold-crystallization transitions were determined from the second heating cycle. Lyophilized nanoparticle formulations were analyzed using the same thermal program to determine whether polymer thermal behavior persisted following nanoparticle processing.

#### *4.5 Dynamic light scattering and zeta-potential analysis*

Nanoparticle hydrodynamic diameter, polydispersity index (PDI), and zeta potential were measured using a Zetasizer Nano ZSP (Malvern Panalytical, Worcestershire, UK). Hydrodynamic diameter and PDI were determined by dynamic light scattering, whereas zeta potential was measured by electrophoretic light scattering. Nanoparticle suspensions were diluted 1:100 in $dH_2O$ before measurement to obtain appropriate scattering intensity and minimize multiple-scattering effects. Hydrodynamic diameter was reported as the intensity-weighted Z-average diameter, PDI was used as a measure of distribution breadth, and zeta potential was determined from electrophoretic mobility. Three (3) independently formulated nanoparticle batches were prepared per group, and then measured in triplicate at 25 °C.

#### *4.6 Scanning electron microscopy*

Nanoparticle morphology was examined by scanning electron microscopy (SEM). Nanoparticle suspensions were prepared at 1 mg $mL^{-1}$, and 2 µL of each suspension was deposited onto an SEM stub and allowed to dry. Dried samples were gold sputter-coated prior to imaging. SEM images were acquired using an Apreo 2S VS instrument at an accelerating voltage of 2.5 kV and a beam current of 0.10 nA. Images were collected at magnifications ranging from 30,000× to 80,000×. The imaging configuration included a pixel width of 4.5 nm, horizontal field width of 13.8 µm, and working distance of 4.0 mm.

SEM images were used to assess nanoparticle morphology and confirm formation of discrete spherical particles

### 5. Acidic incubation and DLS/GPC analysis

Acidic-incubation studies were performed to evaluate degradation-associated colloidal stability and polymer molecular-weight changes under macrophage-relevant acidic conditions. Four nanoparticle formulations were evaluated: $pITA_{C10}^{175\ °C*}$-NPs, $pMSA_{C10}^{200\ °C}$-NPs, $pMSA_{C10}^{175\ °C}$-NPs, and $pMSA_{C10}^{175\ °C*}$-NPs. Lyophilized nanoparticles were dispersed in pH 5.2 acetate buffer at a concentration of 3 mg $mL^{-1}$ and incubated at 37 °C under gentle agitation. Samples were evaluated at 0 h, 24 h, 7 d (168 h), and 14 d (336 h). The 0 h samples were processed immediately following nanoparticle dispersion. At each time point, nanoparticle suspensions were analyzed by DLS to determine hydrodynamic diameter and PDI. These measurements were used to evaluate time-dependent colloidal behavior, including changes consistent with swelling, aggregation, erosion, or particle restructuring.

For GPC analysis, nanoparticles were recovered by centrifugation (12,000 x g, 4 °C, for 10 min), washed twice with deionized water to remove buffer salts, frozen, lyophilized, dissolved in chloroform, filtered, and analyzed using the same GPC method described above. Changes in $M_n$, $M_w$, dispersity, and normalized molecular-weight distributions were evaluated over time.

$M_n$ retention was calculated relative to the formulation-specific recovered-polymer baseline at 0 h according to:

$$M_n = \left(\frac{M_n^t}{M_n^{0h}}\right) \times 100$$

Because the recovered nanoparticle-associated polymer exhibited higher initial $M_n$ values than the corresponding bulk polymers, normalization to the formulation-specific 0 h recovered-polymer value was used to account for potential molecular-weight fractionation during nanoparticle fabrication, purification, and recovery.

The acidic-incubation experiment was interpreted as a material-level measure of acid susceptibility, colloidal restructuring, and degradation-associated molecular-weight change rather than direct evidence of complete intracellular polymer degradation.

### 6. Isolation and differentiation of primary bone marrow-derived macrophages

Male C57BL/6J mice, 5–7 weeks of age, were obtained from The Jackson Laboratory (Bar Harbor, ME, USA) and maintained under pathogen-free conditions at the University of Maryland, Baltimore. All animal procedures were approved by the University of Maryland, Baltimore Institutional Animal Care and Use Committee (IACUC protocol no. 00000975).

Bone marrow was isolated from femurs and tibias and flushed with BMMØ differentiation medium using a 1 mL syringe fitted with a 25-gauge needle. BMMØ medium consisted of RPMI 1640 supplemented with L-glutamine, penicillin (100 units $mL^{-1}$), streptomycin (100 µg $mL^{-1}$), 10 % FBS, and 20 % L929 cell-conditioned medium.

Red blood cells were lysed using ACK lysis buffer. The resulting cell suspension was passed through a 40 µm cell strainer and counted using an EVE automated cell counter with trypan blue exclusion. Cells were plated at $2 \times 10^6$ cells per 10 cm tissue-culture-treated dish and incubated at 37 °C under 5 % $CO_2$. Medium was changed on days 3, 6, and 8. Experiments were performed using differentiated BMMØs on days 7–10.

### 7. Nanoparticle uptake

Macrophages were seeded at 30,000 cells per well in an 8-well Millicell EZ glass chamber slide (MilliporeSigma, Burlington, MA, USA). Cells were treated with 30 µg $mL^{-1}$ Cy5.5-labeled nanoparticles for 24 h. After treatment, cells were washed with 1x HBSS, fixed with fixation buffer, and stained with wheat germ agglutinin (WGA)-Alexa Fluor 594 conjugate (Thermo Fisher Scientific, Waltham, MA, USA). After staining, cells were mounted using Fluoroshield with DAPI (Sigma-Aldrich, St. Louis, MO, USA). Images were taken at 20× magnification using a Nikon Eclipse TI using NIS-Elements 5.42.06 software (Nikon, Melville, NY, USA). Representative images are shown.

### 8. Macrophage viability, apoptosis, and flow-cytometric phenotyping

Bone marrow-derived macrophages were seeded at 0.2 × $10^6$ cells per well in 24-well plates. Where indicated, inflammatory activation was established by overnight stimulation with LPS (100 ng $mL^{-1}$). Cells were subsequently treated for 24 h with pITA- or pMSA-derived nanoparticles (30 µg $mL^{-1}$) or soluble 4OI (125 µM).

#### *8.1 Viability and surface-marker phenotyping*

Flow cytometry staining was performed according to BioLegend protocols with in-house optimization. Cells were first stained with Live/Dead Green Viability/Cytotoxicity dye. Fc receptors were then blocked using anti-CD16/32 antibody (clone 93).

Cells were subsequently stained with fluorophore-conjugated antibodies against macrophage and activation-associated surface markers: PE/Cyanine7 anti-mouse F4/80 (clone BM8), Pacific Blue anti-mouse/human CD11b (clone M1/70), Brilliant Violet 510 anti-mouse CD80 (clone 16-10A1), APC/Cyanine7 anti-mouse CD86 (clone GL-1), and PE-Fire 640 anti-mouse I-A/I-E/MHCII (clone M5/114.15.2). Antibodies were obtained from BioLegend unless otherwise stated.

After surface staining, samples were fixed using BioLegend fixation buffer and acquired using a Cytek Aurora 3 flow cytometer. Data were analyzed using FCS Express 7 (De Novo Software, Glendale, CA, USA). Macrophages were identified based on CD11b and F4/80 expression, following the exclusion of debris and identification of singlets, then CD80, CD86, and MHCII expression was quantified as median fluorescence intensity (MFI), as depicted in the gating strategy **Figure S6**.

#### *8.2 Annexin V/propidium iodide apoptosis analysis*

For apoptosis analysis, cells were blocked with anti-CD16/32 (clone 93), then stained with F4/80 and CD11b followed by FITC-Annexin V and propidium iodide (PI). Samples were kept on ice and analyzed immediately without fixation. Annexin $V^-$/$PI^-$ cells were classified as live, Annexin $V^+$/$PI^-$ cells as early apoptotic, and Annexin $V^+$/$PI^+$ cells as dead/late-stage apoptotic. Data were processed using FCS Express 7.

### 9. RNA sequencing and transcriptomic analysis

RNA isolation, sequencing, and primary data processing were performed by the University of Maryland Institute for Genome Sciences (IGS). Bone marrow-derived macrophages were maintained under resting conditions or stimulated with LPS overnight and treated with pITA-NPs or bioactivity-selected pMSA-NPs at 30 µg $mL^{-1}$ for 24 h. Soluble 4-octyl itaconate (4OI, 125 µM) was included as a soluble immunometabolite comparator. IL-4 (10 ng $mL^{-1}$) stimulated macrophages served as an alternatively activated macrophage control. In this analysis, pMSA-NP refers specifically to the bioactivity-selected $pMSA_{C10}^{200\ °C}$ nanoparticle formulation. After 24 h of treatment, cells were rinsed with PBS, lifted with versine, and transferred to 1.5 mL microcentrifuge tubes. Cells were pelleted and rinsed with sterile PBS using a bench top centrifuge at 500 x g for 5 mins at 4 °C. The pellets were then flash frozen in liquid nitrogen and delivered to IGS for RNA isolation and processing. RNA-seq was performed using n = 3 independent biological replicates per condition.

Gene-level count matrices and DESeq2 differential-expression output files generated by IGS were used for downstream analysis. Differential expression was assessed relative to the appropriate control condition. For resting macrophages, pITA-NP, pMSA-NP, and 4OI groups were compared against untreated resting macrophages. For inflammatory macrophages, LPS + pITA-NP, LPS + pMSA-NP, and LPS + 4OI groups were compared against LPS-treated macrophages. Differentially expressed genes were defined using an absolute $\log_2$ fold change ≥ 0.58 and Benjamini–Hochberg-adjusted P < 0.05.

Downstream analyses were performed in R v4.6.1. Principal component analysis was performed on variance-stabilized expression values. Volcano plots and differentially expressed gene summaries were generated from DESeq2 results. For comparison of pITA-NP and pMSA-NP responses, $\log_2$ fold changes relative to the corresponding M0 or LPS control were compared across genes using Pearson correlation.

Targeted heatmaps were generated from variance-stabilized expression values for predefined genes associated with inflammatory signaling, immunoregulation, immediate-early signaling, metabolic reprogramming, lysosomal/phagocytic function, and oxidative/stress responses, with expression standardized for each gene using row-wise z-score normalization.

Gene set enrichment analysis (GSEA) was performed using ranked DESeq2 results. KEGG enrichment was performed using clusterProfiler with mouse KEGG gene sets following conversion to Entrez gene identifiers, whereas MSigDB Hallmark gene sets were analyzed using clusterProfiler and msigdbr. Enrichment results were summarized using normalized enrichment score (NES) and Benjamini–Hochberg-adjusted false discovery rate (FDR).

**10 LPS-induced endotoxemia survival study**

Male C57BL/6J mice were randomly assigned to treatment groups and administered pITA-NP or bioactivity-selected pMSA-NP by intraperitoneal (i.p.) injection at 2 mg per mouse (10 mg $mL^{-1}$; n = 10 per nanoparticle group) or 200 µL saline vehicle (n = 13). Nanoparticles or vehicle were administered 3 h before i.p. challenge with LPS (20 mg $kg^{-1}$). Animals were monitored for survival for 7 days.

**11. In vivo biodistribution and inflammation study**

A separate cohort of male C57BL/6J mice (6-8 weeks old) was used to evaluate the effects of pITA-NP and bioactivity-selected pMSA-NP during LPS-induced inflammation. Mice received 2 mg $kg^{-1}$ of Cy5.5- labeled pITA-NP or Cy5.5-labeled bioactivity-selected pMSA-NP (5 mice per nanoparticle group) or 200 µL saline for control groups (3 mice per NT and LPS group) i.p. 3 h prior to an i.p.

LPS challenge (5 mg $kg^{-1}$). After 3 h of LPS treatment, the blood plasma and organs were collected for downstream analysis.

#### 11.1 Ex vivo nanoparticle biodistribution

Nanoparticle-associated fluorescence was evaluated ex vivo following systemic administration. After 3 h of LPS exposure, the heart, lungs, spleen, liver, and kidneys were harvested and imaged. Cy5.5 fluorescence intensity within each organ was quantified to compare organ-level nanoparticle-associated signal among treatment groups. Organs were analyzed on the Xenogen in vivo imaging system (IVIS) Spectrum Optical (PerkinElmer, Waltham, MA, USA) with excitation and emission wavelengths of 675 and 720 nm respectively. Representative organ images and quantitative fluorescence measurements were used to assess relative tissue distribution.

#### 11.2 Blood chemistry

Circulating alanine aminotransferase (ALT), aspartate aminotransferase (AST), blood urea nitrogen (BUN), and creatinine were quantified to assess short-term hepatic and renal biochemical responses following treatment. 3 h post-LPS exposure, blood was extracted and collected into EDTA coated collection tube to prevent clotting. Samples were centrifuged at 1500 x g for 10 min at 4 °C. The plasma was then collected, frozen, then sent for analysis by VRL Diagnostics (Gaithersburg, MD, USA).

#### 11.3 Multiplex cytokine and chemokine analysis

Circulating cytokines and chemokines were quantified using a 26-plex Luminex panel (EPXR260-26088-901) and processed using the Luminex xPONENT® software (Thermo Fisher Scientific, Waltham, MA, USA), following the manufacturer's protocol. Concentrations were determined for individual animals and retained as absolute concentration values for the corresponding Supporting Information analyses. For normalized analyses shown in the main manuscript, each analyte was normalized relative to the geometric mean concentration of the corresponding LPS control group and expressed as $\log_2$ fold change. Thus, individual LPS-control values were retained rather than artificially set to zero; only the geometric mean of the LPS group corresponds to a normalized $\log_2$ fold change of zero.

#### 11.4 H&E histology staining

Organ damage and alterations in tissue-level architecture was assessed via histopathological evaluation. Following IVIS, organs were rinsed in 1x PBS, placed into tissue cassettes, and fixed for 24 h in 10 % buffered formalin (Sigma-Aldrich, St. Louis, MO, USA). The following day, samples were rinsed with PBS and delivered to the Pathology Biorepository Shared Services Core at the University of Maryland, Baltimore, where the samples were then paraffin embedded and sectioned into 5 µm slices. Slides were stained with hematoxylin and eosin (H&E). Images were taken at 4× and 10× magnification using an ECHO revolve microscope (Discover Echo Inc., San Diego, CA, USA). Representative images shown.

### 12. Statistical Analysis

Statistical analyses were performed using GraphPad Prism v11.1.0 (GraphPad Software, Boston, MA, USA), except for transcriptomic analyses performed in R as described above. Unless otherwise indicated, data are presented as mean ± standard deviation (SD), with individual independent data points shown where applicable.

For macrophage viability, apoptosis, and flow-cytometric phenotyping experiments, comparisons among multiple treatment groups were performed using one-way analysis of variance (ANOVA) followed by Dunnett's multiple-comparisons test against the corresponding control group. Selected pairwise comparisons were analyzed using two-sided unpaired t-tests with Welch's correction. Exact P values are reported where applicable.

For ex vivo biodistribution measurements, data were analyzed using two-way ANOVA followed by Tukey's multiple-comparisons test. Circulating cytokine and chemokine data were analyzed using the Kruskal–Wallis test followed by Dunn's multiple-comparisons test against the LPS control.

Survival distributions were compared using the log-rank (Mantel–Cox) test. Each nanoparticle-treated survival curve was compared with the vehicle-control curve, and multiplicity was controlled using the Holm–Šídák method. Multiplicity-adjusted P values were reported using a familywise $\alpha$ threshold of 0.05.

For RNA-seq analyses, differentially expressed genes were identified using DESeq2 with $|\log_2$ fold change$| \geq 0.58$ and Benjamini–Hochberg-adjusted $P < 0.05$. Pathway-enrichment analyses were summarized using normalized enrichment scores and Benjamini–Hochberg-adjusted false discovery rates as described above. Statistical significance was defined as $P < 0.05$ or adjusted $P < 0.05$, as appropriate.

The definition of n was experiment-specific and is provided in the corresponding figure or table legend. Polymer-synthesis and polymerization-kinetics data were obtained from three independent synthesis batches (n = 3). For acidic-incubation GPC and DLS analyses, n = 3 measurements per formulation and time point. Cell-based studies used independent biological experiments, RNA-seq used n = 3 independent biological replicates per condition, and each n in the *in vivo* studies represents an individual animal.

**Table S1. Reaction conditions used for $pITA_{C10}$ and $pMSA_{C10}$ synthesis**

<table>
<tr><th>Polymer candidate</th><th>Acid monomer</th><th>Acid amount</th><th>1,10-Decanediol</th><th>$SnCl_2$</th><th>MeHQ</th><th>Temperature</th><th>Reaction atmosphere</th></tr>
<tr><td>$pITA_{C10}^{175\ °C*}$</td><td>Itaconic acid</td><td rowspan="4">650 mg, 5.0 mmol</td><td rowspan="4">871.5 mg, 5.0 mmol</td><td rowspan="4">107 mg, 0.56 mmol</td><td>7 mg, 0.056 mmol</td><td>175 °C</td><td rowspan="4">1 h Ar<br>+<br>1 h vacuum</td></tr>
<tr><td>$pMSA_{C10}^{200\ °C}$</td><td rowspan="3">Mesaconic acid</td><td>-</td><td>200 °C</td></tr>
<tr><td>$pMSA_{C10}^{175\ °C}$</td><td>-</td><td>175 °C</td></tr>
<tr><td>$pMSA_{C10}^{175\ °C*}$</td><td>7 mg, 0.056 mmol</td><td>175 °C</td></tr>
</table>

**Table S2. Molecular-weight characteristics of synthesized itaconate- and mesaconate-based polyesters.**

| Polymer Formulation | Synthesis Temperature (°C) | Inhibitor (MeHQ) | $M_n$ (kDa) | Đ ($M_w/M_n$) |
|---|---|---|---|---|
| $pITA_{C10}$ | 175 | Yes | 27.92 ± 2.31 | 2.27 ± 0.31 |
| $pMSA_{C10}$ | 200 | No | 8.62 ± 0.88 | 1.90 ± 0.20 |
| $pMSA_{C10}$ | 175 | Yes | 9.82 ± 0.14 | 1.57 ± 0.21 |
| $pMSA_{C10}$ | 175 | No | 8.54 ± 0.65 | 1.77 ± 0.15 |

*Values are presented as mean ± SD (n = 3). $M_n$, number-average molecular weight; Đ, dispersity ($M_w/M_n$); MeHQ, 4-methoxyphenol.*

**Table S3. Evolution of molecular weight and dispersity during the synthesis of itaconate- and mesaconate-based polyesters.**

| Polymer formulation | Polymerization time (h) | $M_n$ (kDa) | Đ ($M_w/M_n$) |
|---|---|---|---|
| $pITA_{C10}^{175\ °C*}$ | 0.5 | 4.54 ± 1.06 | 1.20 ± 0.10 |
| | 1.0 | 18.79 ± 1.51 | 1.57 ± 0.15 |
| | 1.5 | 26.53 ± 2.64 | 1.80 ± 0.26 |
| | 2.0 | 28.58 ± 3.41 | 2.17 ± 0.31 |
| | 2.5 | 29.89 ± 4.07 | 2.33 ± 0.32 |
| $pMSA_{C10}^{200\ °C}$ | 0.5 | 5.63 ± 1.47 | 1.37 ± 0.21 |
| | 1.0 | 7.83 ± 0.83 | 1.60 ± 0.17 |
| | 1.5 | 8.21 ± 1.03 | 1.70 ± 0.20 |
| | 2.0 | 8.47 ± 1.09 | 1.77 ± 0.35 |
| | 2.5 | 8.62 ± 1.05 | 1.87 ± 0.31 |
| $pMSA_{C10}^{175\ °C*}$ | 0.5 | 4.96 ± 0.68 | 1.27 ± 0.21 |
| | 1.0 | 7.79 ± 1.13 | 1.33 ± 0.25 |
| | 1.5 | 8.86 ± 0.66 | 1.47 ± 0.21 |
| | 2.0 | 9.54 ± 0.69 | 1.60 ± 0.20 |
| | 2.5 | 9.86 ± 0.70 | 1.63 ± 0.21 |
| $pMSA_{C10}^{175\ °C}$ | 0.5 | 4.47 ± 0.44 | 1.23 ± 0.15 |
| | 1.0 | 6.38 ± 0.57 | 1.50 ± 0.26 |
| | 1.5 | 7.64 ± 1.08 | 1.63 ± 0.15 |
| | 2.0 | 8.16 ± 0.85 | 1.60 ± 0.10 |
| | 2.5 | 8.51 ± 0.71 | 1.70 ± 0.10 |

*Values are presented as mean ± SD (n = 3). $M_n$, number-average molecular weight; Đ, dispersity ($M_w/M_n$).*

**Table S4. Molecular-weight and dispersity changes in polymer fractions recovered from itaconate- and mesaconate-derived nanoparticles during acidic incubation.**

| Polymer formulation | Incubation time (h) | $M_n$ (kDa) | Đ ($M_w/M_n$) |
|---|---|---|---|
| $pITA_{C10}^{175\ °C*}$ | 0 | 35.40 ± 3.73 | 3.57 ± 0.29 |
| | 24 | 32.06 ± 3.79 | 4.79 ± 0.81 |
| | 168 | 26.93 ± 4.28 | 3.76 ± 0.33 |
| | 336 | 14.62 ± 0.74 | 3.77 ± 0.97 |
| $pMSA_{C10}^{200\ °C}$ | 0 | 16.49 ± 1.86 | 1.46 ± 0.11 |
| | 24 | 13.28 ± 2.62 | 1.40 ± 0.21 |
| | 168 | 9.19 ± 1.21 | 1.70 ± 0.52 |
| | 336 | 6.04 ± 1.55 | 1.57 ± 0.17 |
| $pMSA_{C10}^{175\ °C*}$ | 0 | 16.25 ± 1.00 | 1.57 ± 0.39 |
| | 24 | 14.05 ± 0.67 | 2.17 ± 0.64 |
| | 168 | 10.79 ± 0.98 | 1.99 ± 0.51 |
| | 336 | 8.48 ± 0.50 | 2.05 ± 0.20 |
| $pMSA_{C10}^{175\ °C}$ | 0 | 17.01 ± 1.63 | 1.67 ± 0.23 |
| | 24 | 15.65 ± 0.75 | 1.75 ± 0.23 |
| | 168 | 13.03 ± 0.54 | 2.20 ± 0.25 |
| | 336 | 11.44 ± 0.71 | 1.98 ± 0.27 |

*Values are presented as mean ± SD (n = 3 measurements per formulation and time point). $M_n$, number-average molecular weight; Đ, dispersity ($M_w/M_n$). Nanoparticles were incubated in pH 5.2 acetate buffer at 37 ℃. The 0 h values represent the polymer fraction recovered after brief exposure to acidic buffer and immediate washing and recovery.*

**Table S5. Antibodies and staining reagents used for flow-cytometric analysis of BMMØs**

| Marker | Fluorophore | Clone | Supplier | Purpose |
|---|---|---|---|---|
| F4/80 | PE/Cyanine7 | BM8 | BioLegend | Macrophage identification |
| CD11b | Pacific Blue | M1/70 | BioLegend | Myeloid/macrophage identification |
| CD80 | Brilliant Violet 510 | 16-10A1 | BioLegend | Costimulatory/activation-associate marker |
| CD86 | APC/Cyanine7 | GL-1 | BioLegend | Costimulatory/activation-associated marker |
| MHCII, I-A/I-E | PE-Fire 640 | M5/114.15.2 | BioLegend | Antigen-presentation-associated marker |
| CD16/32 | - | 93 | BioLegend | Fc receptor blocking |
| Live/Dead Green | - | - | Thermo Fisher Scientific | Viability assessment |

**Table S6. Summary of differentially expressed genes following various treatments under M0 or pre-established LPS-activated macrophage polarization conditions.**

| Treatment condition | DEG comparison | Direction | # genes |
|---|---|---|---|
| M0 | 4OI vs NT | Downregulated | 2535 |
| M0 | 4OI vs NT | Upregulated | 1687 |
| M0 | pITA-NP vs NT | Downregulated | 37 |
| M0 | pITA-NP vs NT | Upregulated | 34 |
| M0 | pMSA-NP vs NT | Downregulated | 181 |
| M0 | pMSA-NP vs NT | Upregulated | 123 |
| LPS-activated | LPS + 4OI vs LPS | Downregulated | 1882 |
| LPS-activated | LPS + 4OI vs LPS | Upregulated | 1010 |
| LPS-activated | LPS + pITA-NP vs LPS | Downregulated | 12 |
| LPS-activated | LPS + pITA-NP vs LPS | Upregulated | 68 |
| LPS-activated | LPS + pMSA-NP vs LPS | Downregulated | 23 |
| LPS-activated | LPS + pMSA-NP vs LPS | Upregulated | 51 |

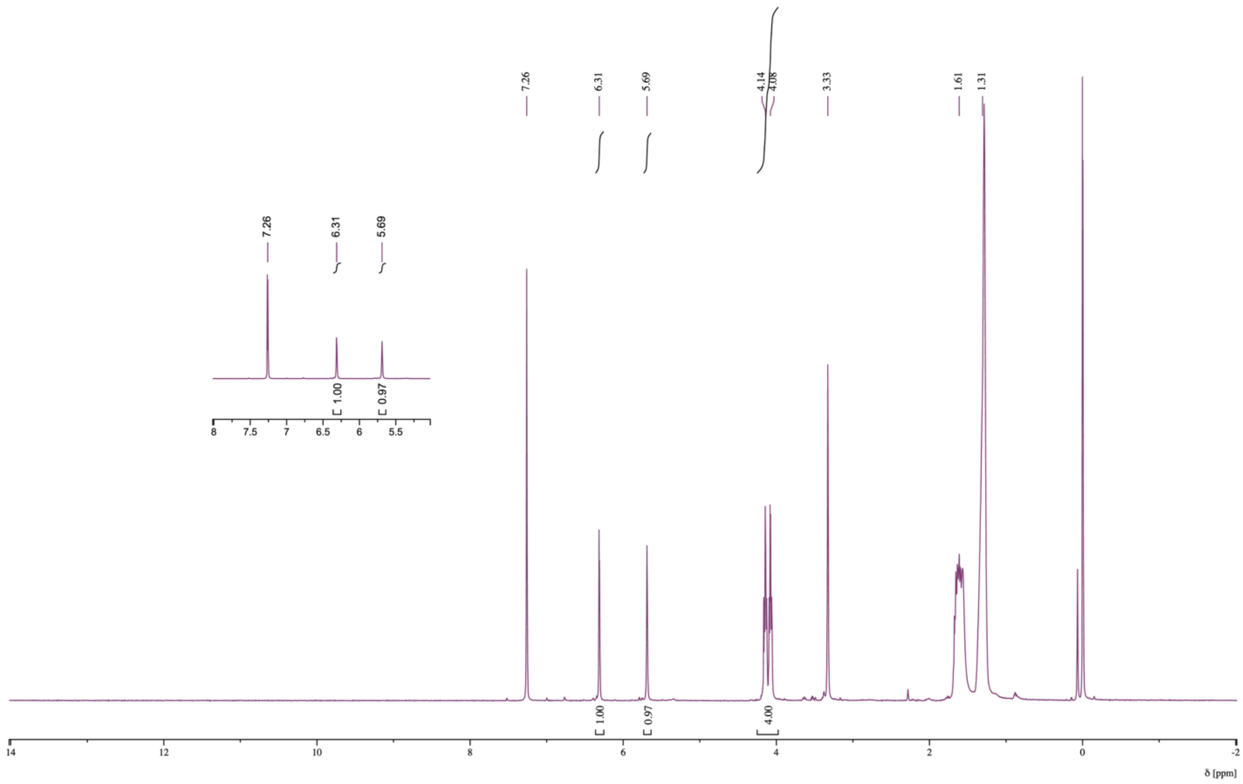


**Figure S1. $^1$H NMR spectrum of pITA$_{C10}$ synthesized at 175 °C with MeHQ.** The inset highlights the characteristic itaconate-derived exo-methylene resonances. $^1$H NMR (400 MHz, $CDCl_3$, δ ppm): 6.31 and 5.69 (itaconate exo-methylene, $=CH_2$, 2H), 4.14–4.08 (ester-adjacent $–OCH_2–$, 4H), 3.33 (itaconate backbone $–CH_2–$, 2H), 1.61 ($–OCH_2–CH_2–$, 4H), 1.31 (internal decanediol $–CH_2–$, 12H).

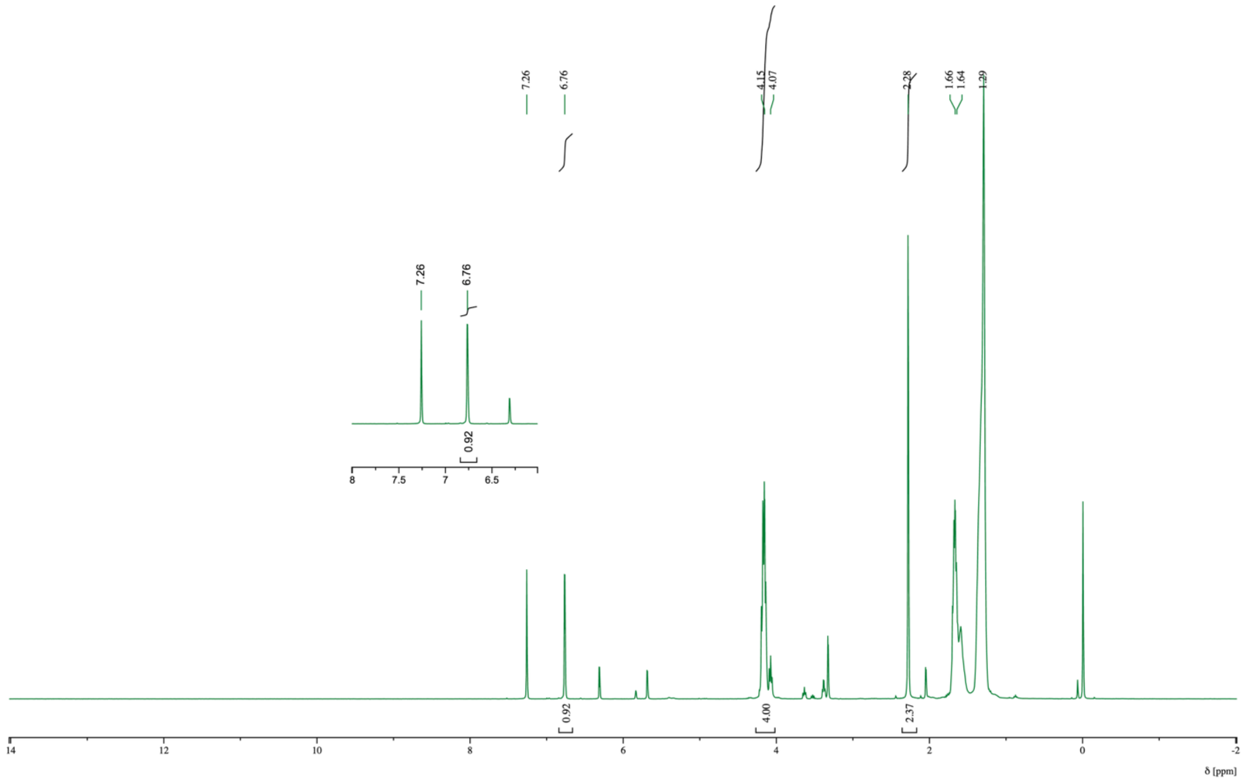


**Figure S2. $^1$H NMR spectrum of pMSA$_{C10}$ synthesized at 200 °C without MeHQ.** The inset highlights the characteristic mesaconate-derived internal alkene resonance. $^1$H NMR (400 MHz, $CDCl_3$, δ ppm): 6.76 (mesaconate internal alkene, –CH=C($CH_3$)–, 1H), 4.15–4.07 (ester-adjacent –$OCH_2$–, 4H), 2.26 (vinylic methyl, –C($CH_3$)=, 3H), 1.66–1.64 (–$OCH_2$–$CH_2$–, 4H), 1.29 (internal decanediol –$CH_2$–, 12H).

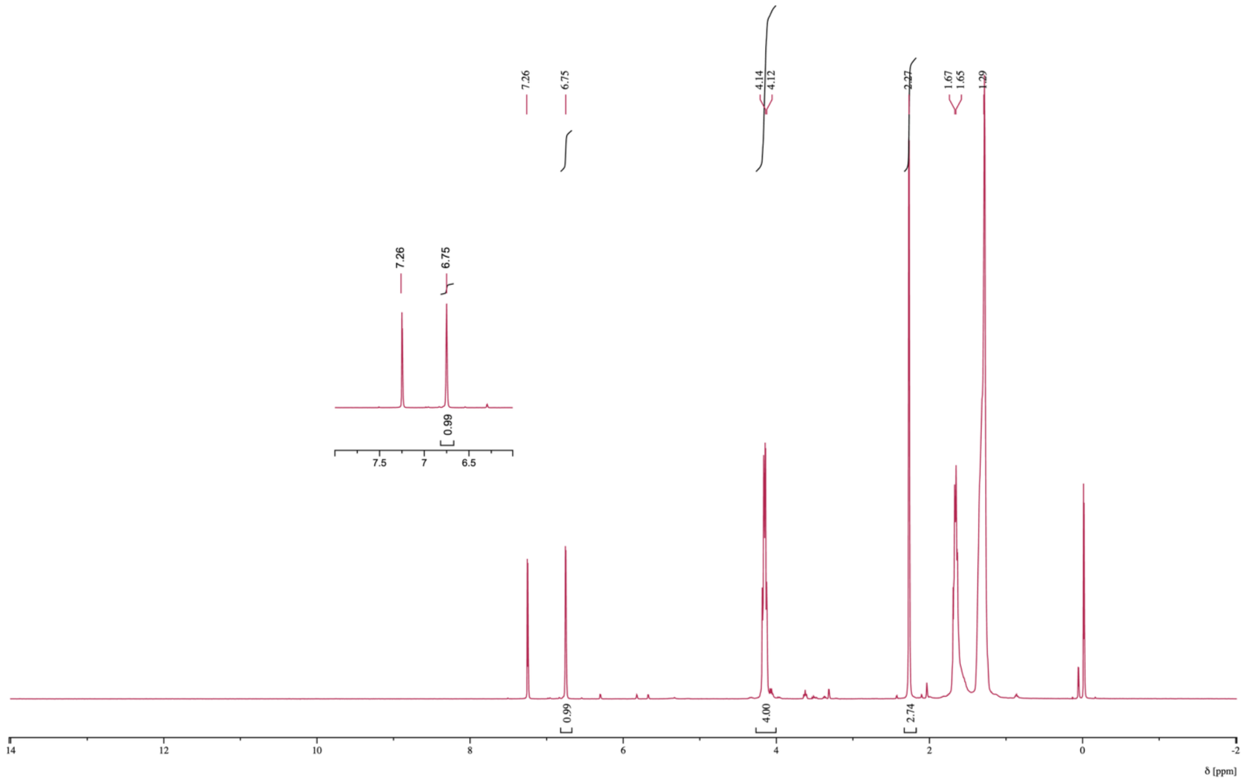


**Figure S3. $^1$H NMR spectrum of pMSA$_{C10}$ synthesized at 175 °C without MeHQ.** The inset highlights the characteristic mesaconate-derived internal alkene resonance. $^1$H NMR (400 MHz, $CDCl_3$, δ ppm): 6.75 (mesaconate internal alkene, –CH=C($CH_3$)–, 1H), 4.14–4.12 (ester-adjacent –$OCH_2$–, 4H), 2.27 (vinylic methyl, –C($CH_3$)=, 3H), 1.67–1.65 (–$OCH_2$–$CH_2$–, 4H), 1.29 (internal decanediol –$CH_2$–, 12H).

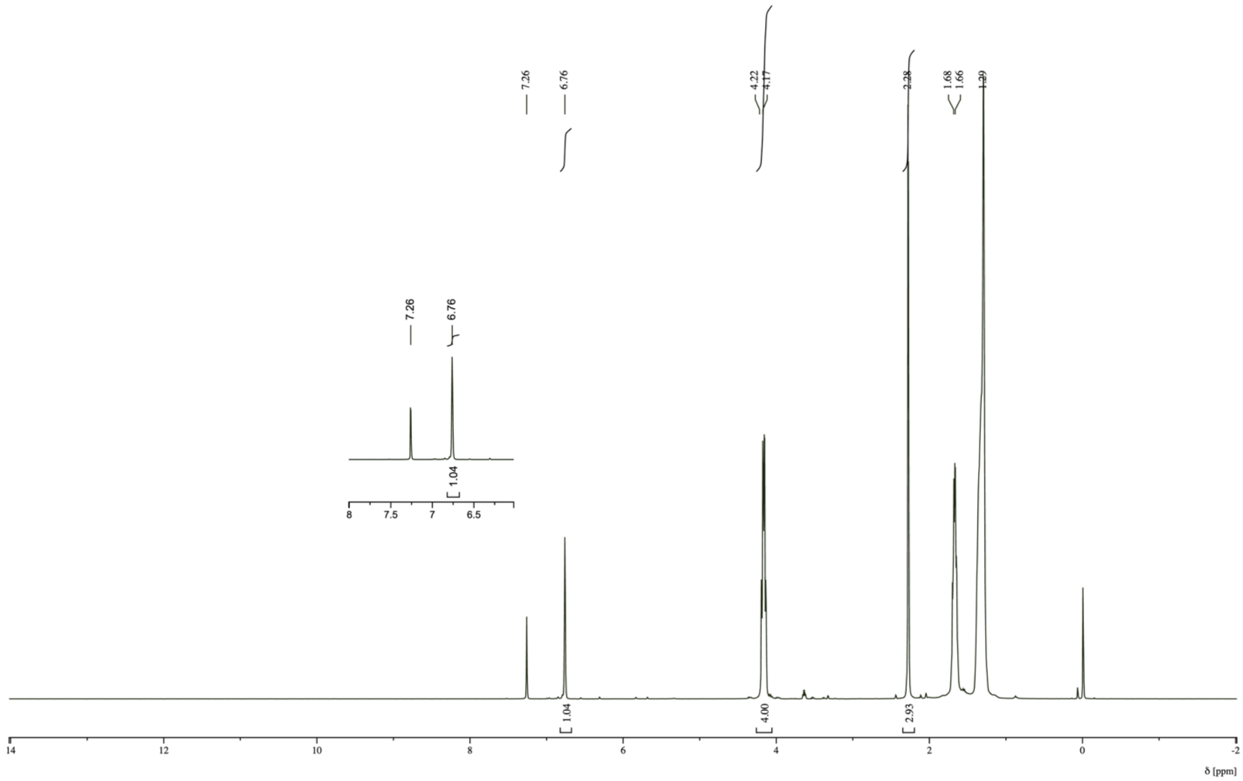


**Figure S4. [1]H NMR spectrum of pMSA$_{C10}$ synthesized at 175 °C with MeHQ.** The inset highlights the characteristic mesaconate-derived internal alkene resonance. $^1$H NMR (400 MHz, $CDCl_3$, δ ppm): 6.76 (mesaconate internal alkene, –CH=C($CH_3$)–, 1H), 4.22–4.17 (ester-adjacent –$OCH_2$–, 4H), 2.28 (vinylic methyl, –C($CH_3$)=, 3H), 1.68–1.66 (–$OCH_2$–$CH_2$–, 4H), 1.29 (internal decanediol –$CH_2$–, 12H).

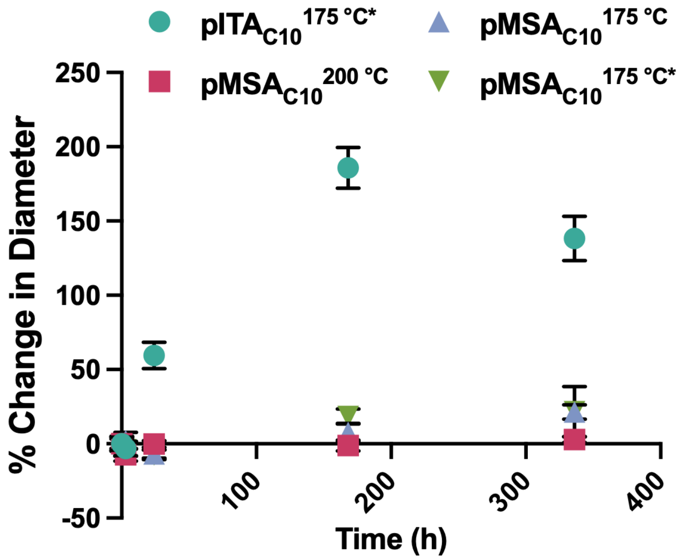


**Figure S5. Acidic incubation–associated changes in nanoparticle hydrodynamic diameter.** Percentage change in hydrodynamic diameter of pITA$_{C10}$$^{175\ °C*}$-NPs and pMSA$_{C10}$ formulations during incubation in pH 5.2 acetate buffer at 37 °C, measured by DLS at 0, 24, 168, and 336 h. Data are mean ± SD (n = 3 independent nanoparticle batches).

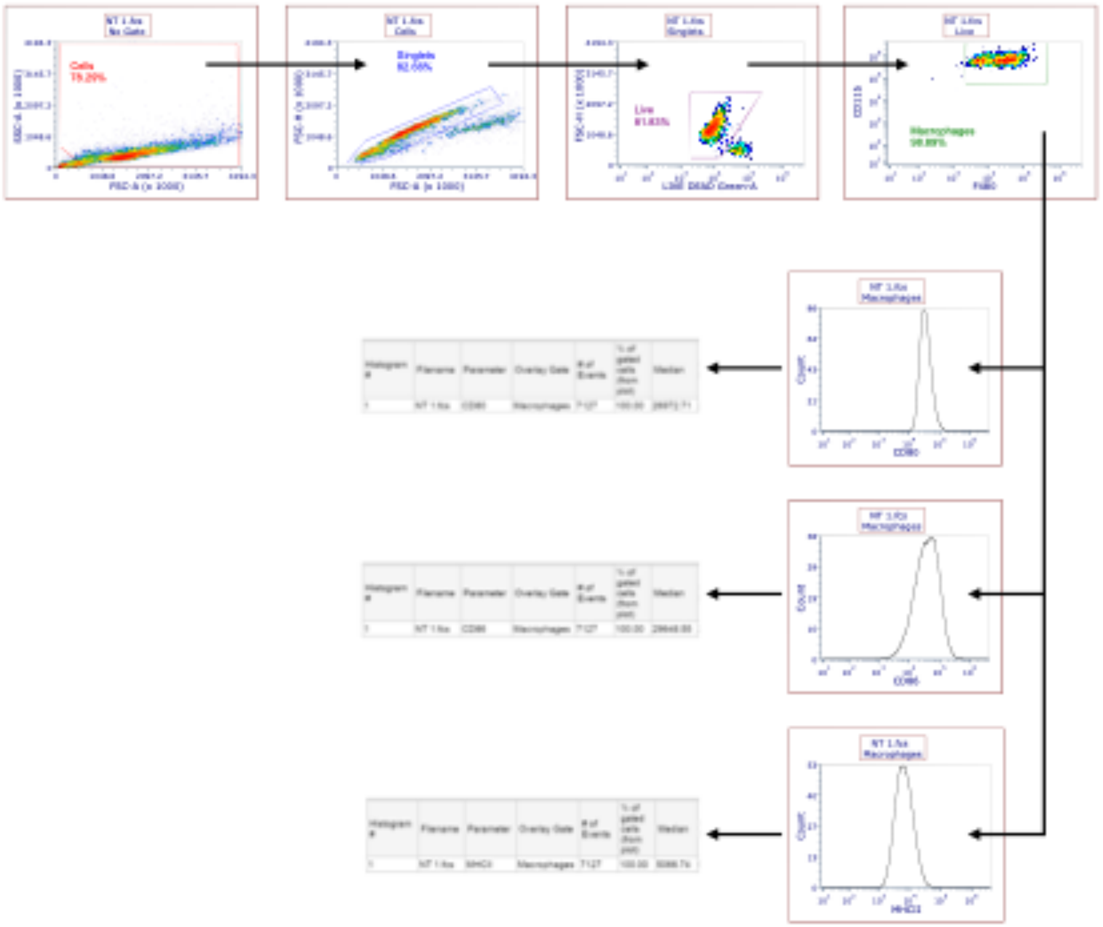

**Figure S6. Representative gating strategy for macrophage phenotype.** Macrophages were stained, fixed, then analyzed on a Cytek Aurora 3 flow cytometer and gated as shown. The median value of the histogram surface markers was used as the MFI for data presentation.

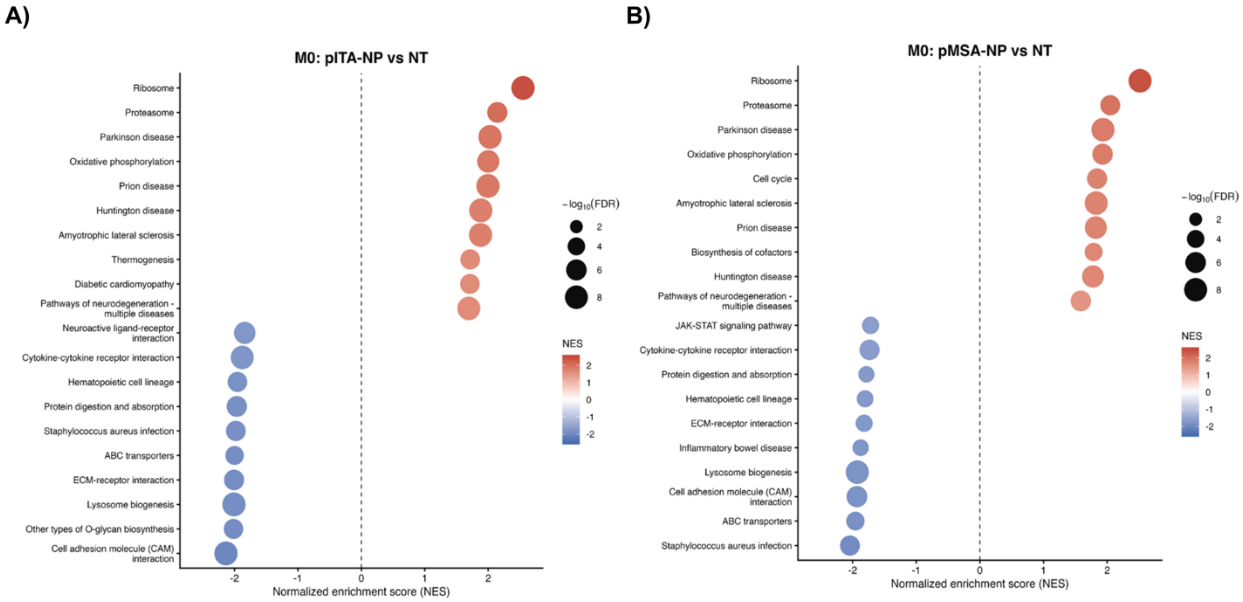


**Figure S7. KEGG gene set enrichment analysis of pITA-NP- and pMSA-NP-treated M0 macrophages.** Gene set enrichment analysis (GSEA) of KEGG pathways following treatment of M0 bone marrow-derived macrophages with **(A)** pITA-NP or **(B)** pMSA-NP relative to untreated (NT) controls. Pathways with the strongest positive and negative enrichment are shown. Dot position and color indicate the normalized enrichment score (NES), with positive and negative values representing positive and negative enrichment, respectively, and dot size represents $-\log_{10}$(FDR). RNA-seq was performed using n = 3 biological replicates per condition.

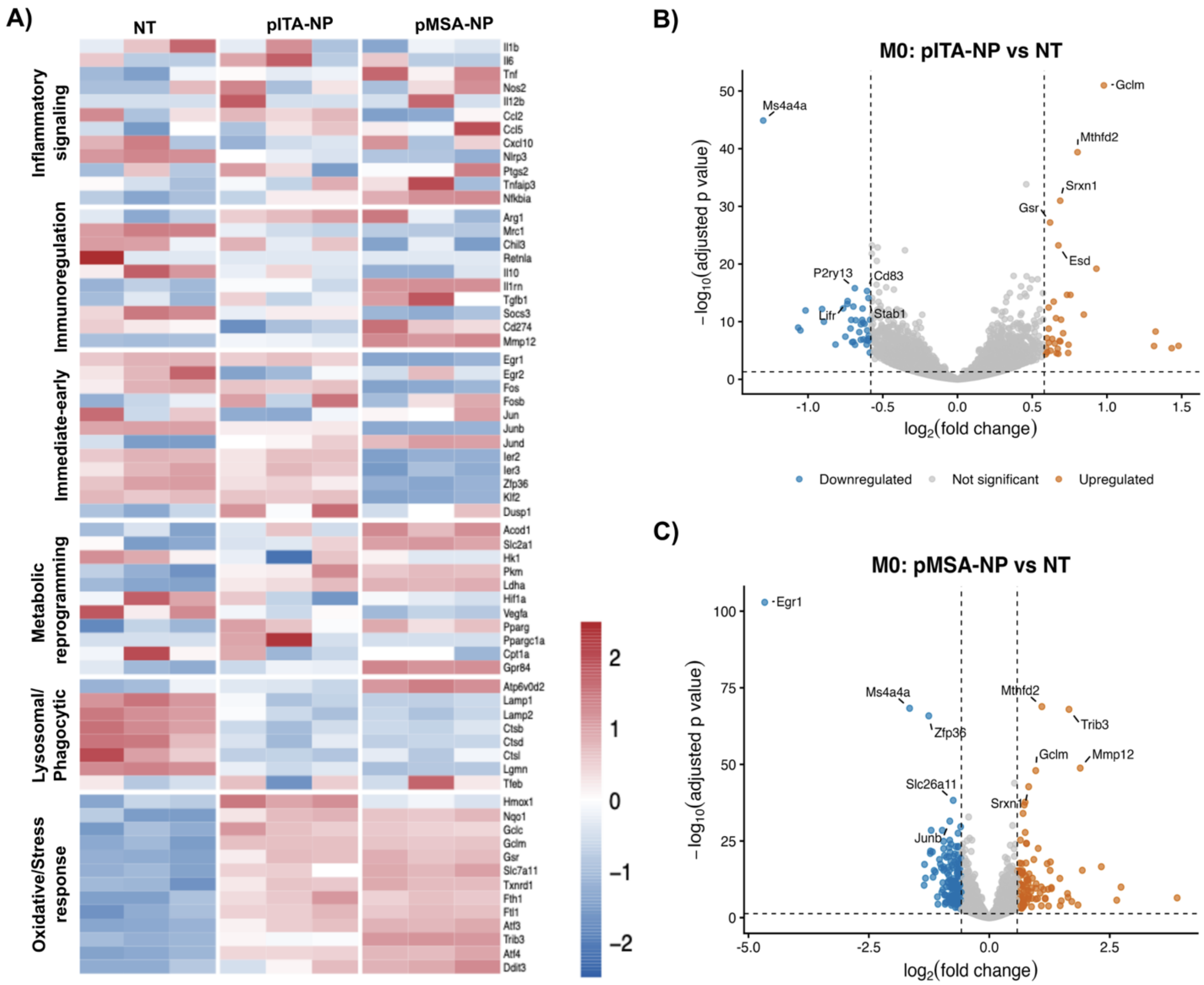


**Figure S8. Gene-level transcriptional responses to pITA-NP and pMSA-NP treatment in M0 macrophages. (A)** Targeted heatmap showing expression patterns of genes grouped into inflammatory signaling, immunoregulation, immediate-early signaling, metabolic reprogramming, lysosomal/phagocytic function, and oxidative/stress-response programs in untreated (NT), pITA-NP-treated, and pMSA-NP-treated M0 macrophages. Expression values are shown as row-scaled relative expression across biological replicates. **(B,C)** Volcano plots showing differentially expressed genes following **(B)** pITA-NP or **(C)** pMSA-NP treatment relative to NT controls. Genes meeting the differential-expression threshold (adjusted $p < 0.05$ and $|\log_2$ fold change$| \geq 0.58$) are shown as upregulated or downregulated, with selected genes labeled. RNA-seq was performed using n = 3 biological replicates per condition.

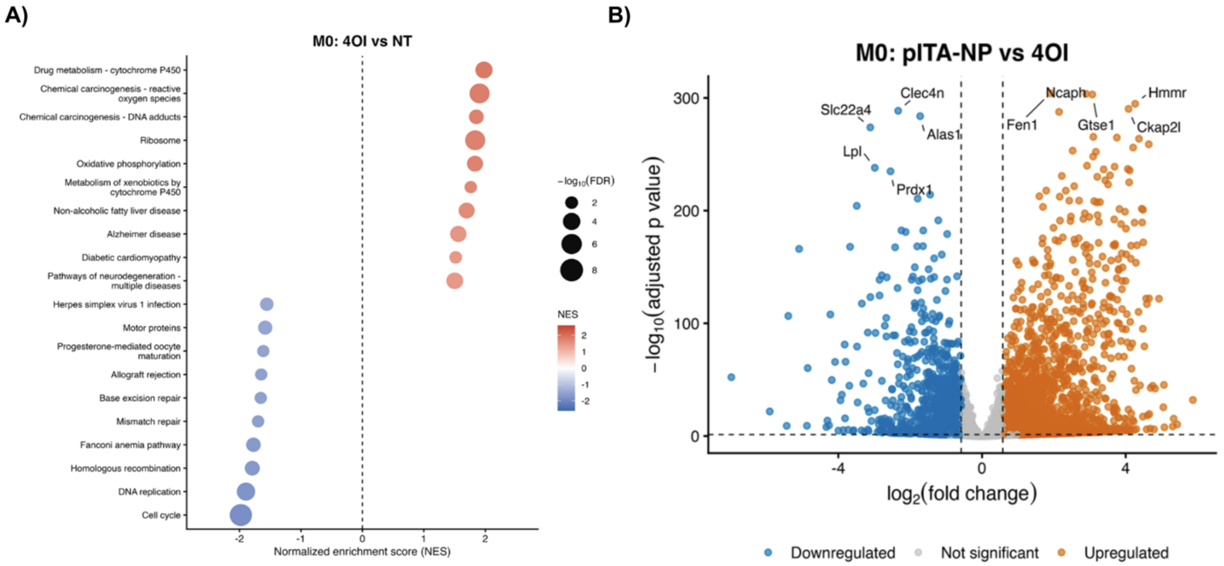


**Figure S9. Transcriptional comparison of 4OI and pITA-NP in M0 macrophages. (A)** Gene set enrichment analysis (GSEA) of KEGG pathways following 4OI treatment relative to untreated (NT) M0 macrophages. Dot position and color indicate the normalized enrichment score (NES), with positive and negative values representing positive and negative enrichment, respectively, and dot size represents $-\log_{10}$(FDR). **(B)** Volcano plot showing differential gene expression between pITA-NP- and 4OI-treated M0 macrophages. Genes meeting the differential-expression threshold (adjusted $p < 0.05$ and $|\log_2$ fold change$| \geq 0.58$) are shown as upregulated or downregulated, with selected genes labeled. RNA-seq was performed using $n = 3$ biological replicates per condition.

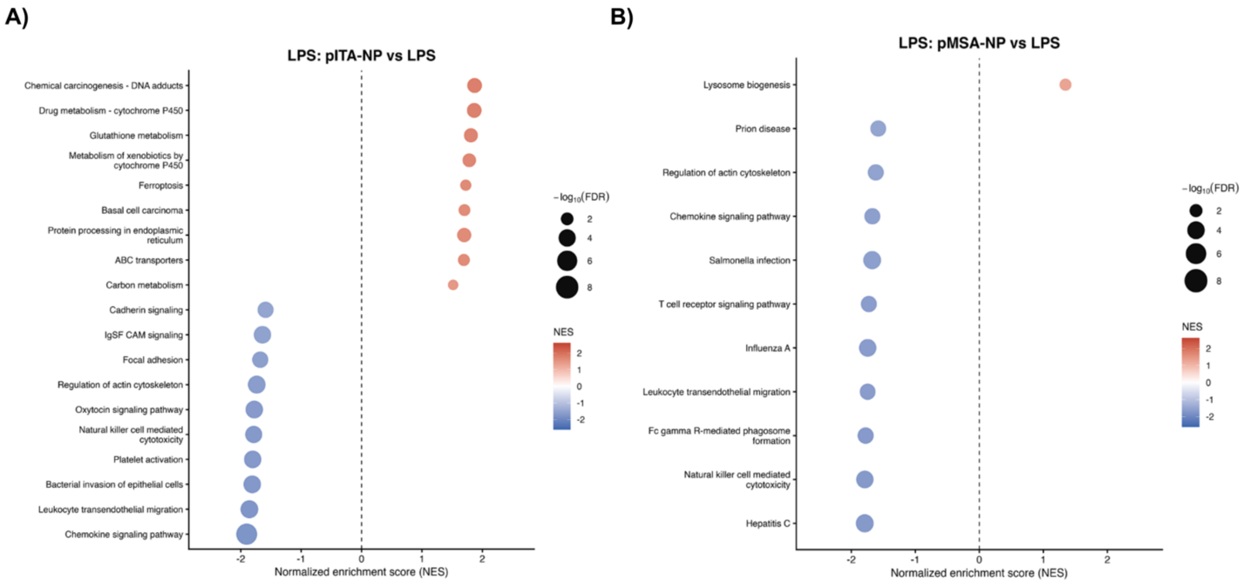


**Figure S10. KEGG gene set enrichment analysis of pITA-NP- and pMSA-NP-treated LPS-activated macrophages.** Gene set enrichment analysis (GSEA) of KEGG pathways following treatment of LPS-activated bone marrow-derived macrophages with **(A)** pITA-NP or **(B)** pMSA-NP relative to LPS controls. Pathways with the strongest positive and negative enrichment are shown. Dot position and color indicate the normalized enrichment score (NES), with positive and negative values representing positive and negative enrichment, respectively, and dot size represents $-\log_{10}$(FDR). RNA-seq was performed using n = 3 biological replicates per condition.

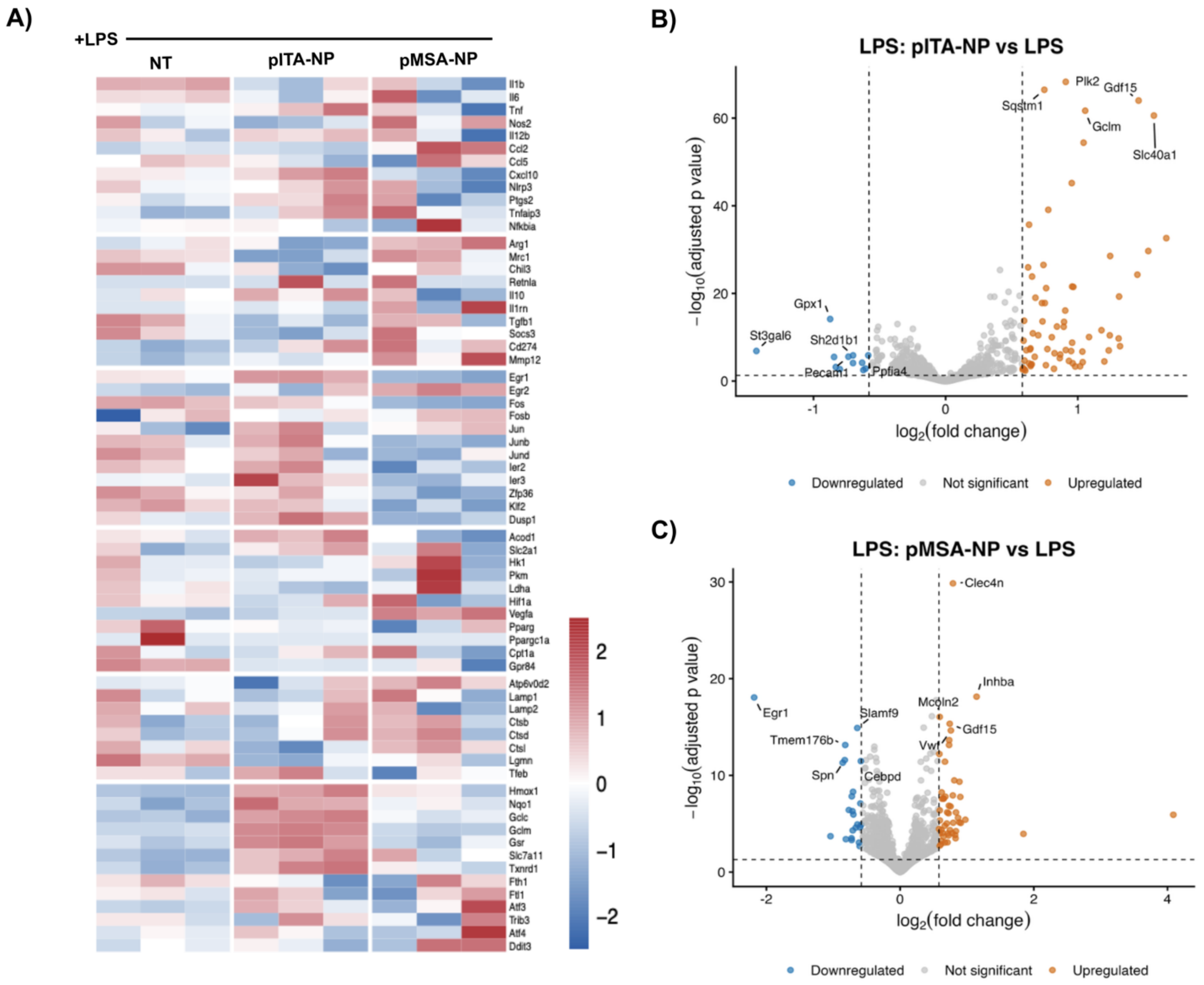


**Figure S11. Gene-level transcriptional responses to pITA-NP and pMSA-NP treatment in LPS-activated macrophages. (A)** Targeted heatmap showing expression patterns of genes grouped into inflammatory signaling, immunoregulation, immediate-early signaling, metabolic reprogramming, lysosomal/phagocytic function, and oxidative/stress-response programs in LPS-activated macrophages treated with pITA-NP or pMSA-NP. Expression values are shown as row-scaled relative expression across biological replicates. **(B,C)** Volcano plots showing differentially expressed genes following **(B)** pITA-NP or **(C)** pMSA-NP treatment relative to LPS controls. Genes meeting the differential-expression threshold (adjusted $p < 0.05$ and $|\log_2 \text{fold change}| \geq 0.58$) are shown as upregulated or downregulated, with selected genes labeled. RNA-seq was performed using n = 3 biological replicates per condition.

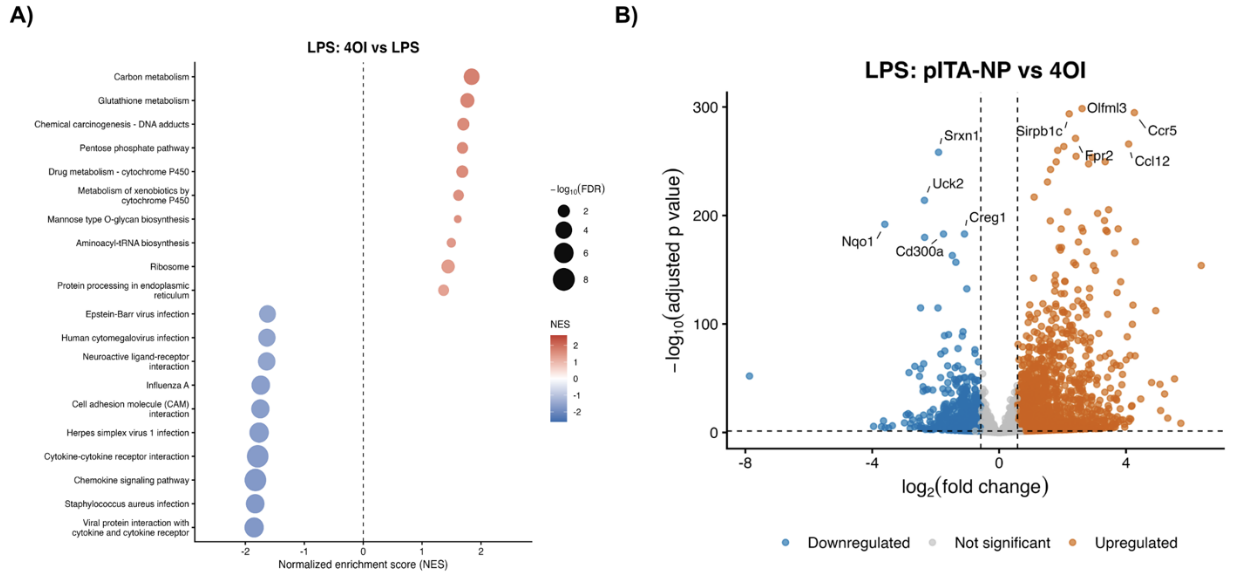


**Figure S12. Transcriptional comparison of 4OI and pITA-NP in LPS-activated macrophages. (A)** Gene set enrichment analysis (GSEA) of KEGG pathways following 4OI treatment relative to LPS-activated macrophages. Dot position and color indicate the normalized enrichment score (NES), with positive and negative values representing positive and negative enrichment, respectively, and dot size represents $-\log_{10}$(FDR). **(B)** Volcano plot showing differential gene expression between pITA-NP- and 4OI-treated LPS-activated macrophages. Positive $\log_2$ fold-change values indicate higher expression following pITA-NP treatment, whereas negative values indicate higher expression following 4OI treatment. Genes meeting the differential-expression threshold (adjusted $p < 0.05$ and $|\log_2$ fold change$| \geq 0.58$) are shown as upregulated or downregulated, with selected genes labeled. RNA-seq was performed using $n = 3$ biological replicates per condition.

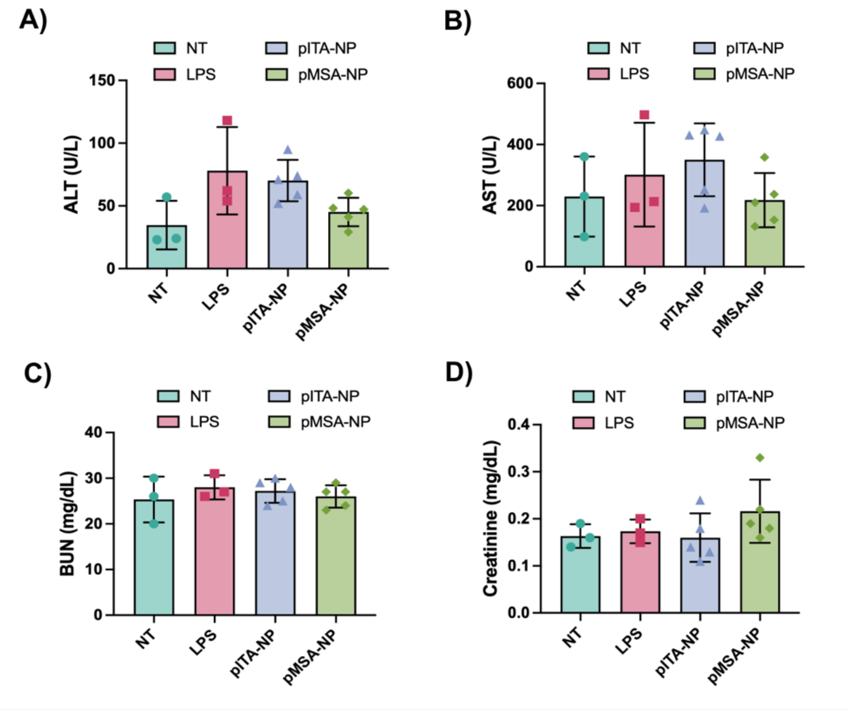


**Figure S13. Serum biochemical markers following metabolite-derived nanoparticle pretreatment and LPS challenge. (A)** Alanine aminotransferase (ALT), **(B)** aspartate aminotransferase (AST), **(C)** blood urea nitrogen (BUN), and **(D)** creatinine levels in NT, LPS, pITA-NP, and pMSA-NP groups. ALT and AST were assessed as hepatic biochemical markers, whereas BUN and creatinine were assessed as renal biochemical markers. Data are presented as mean ± SD, with individual symbols representing individual animals. Statistical comparisons were performed using one-way ANOVA followed by Dunnett's multiple-comparisons test against the LPS group.

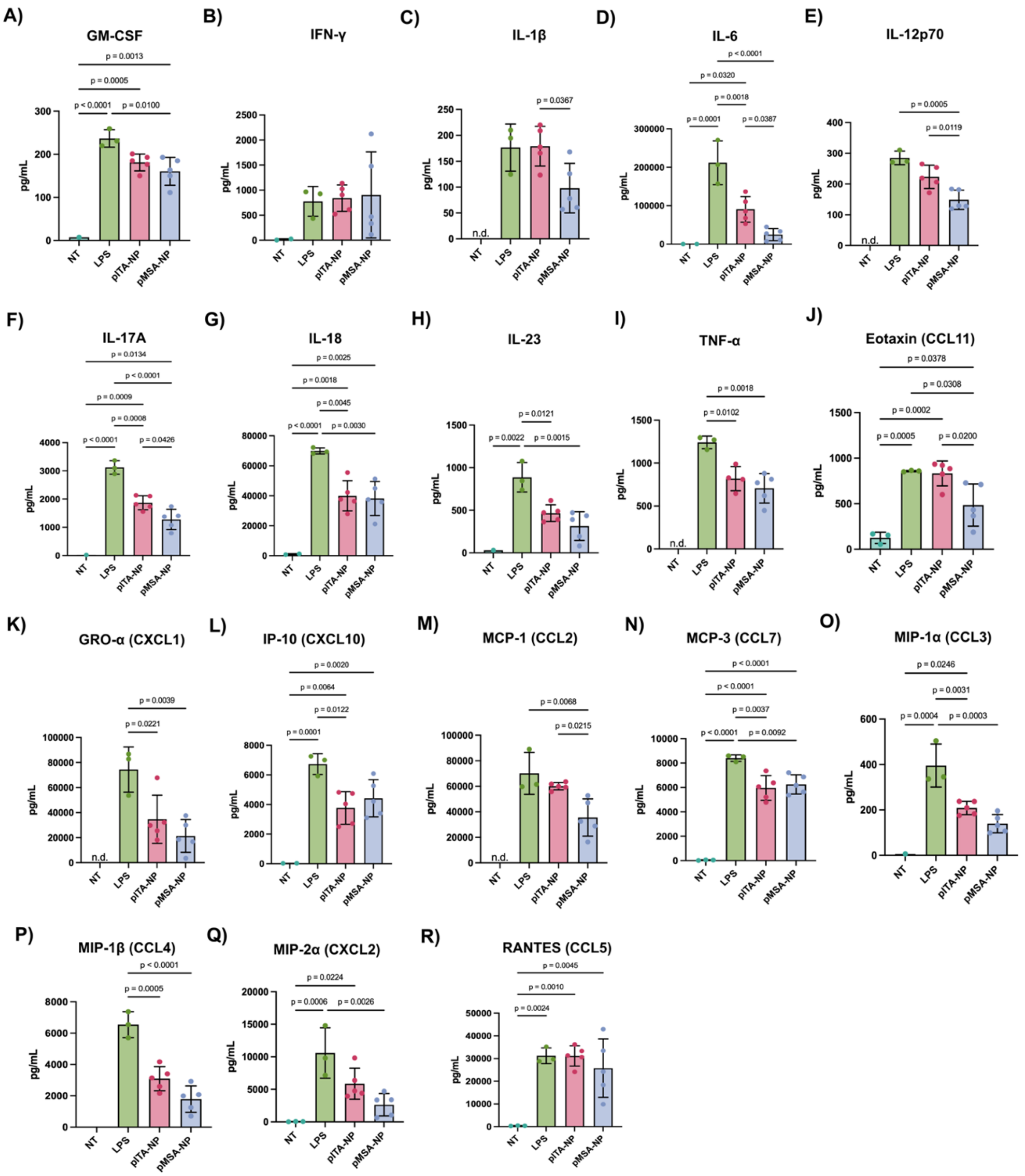


**Figure S14. Circulating inflammatory cytokine and chemokine responses following metabolite-derived nanoparticle pretreatment in LPS-induced endotoxemia. Concentrations of (A)** GM-CSF, **(B)** IFN-γ, **(C)** IL-1β, **(D)** IL-6, **(E)** IL-12p70, **(F)** IL-17A, **(G)** IL-18, **(H)** IL-23, **(I)** TNF-α, **(J)** eotaxin/CCL11, **(K)** GRO-α/CXCL1, **(L)** IP-10/CXCL10, **(M)** MCP-1/CCL2, **(N)** MCP-3/CCL7, **(O)** MIP-1α/CCL3, **(P)** MIP-1β/CCL4, **(Q)** MIP-2α/CXCL2, and **(R)** RANTES/CCL5 in untreated (NT), LPS-treated, pITA-NP-treated, and pMSA-NP-treated mice. Data are presented as mean ± SD, with individual points representing individual animals. Statistical comparisons were performed using the Kruskal–Wallis test followed by Dunn's multiple-comparisons test; exact adjusted P values are shown for the indicated comparisons. Cytokine concentrations are reported in pg $mL^{-1}$.

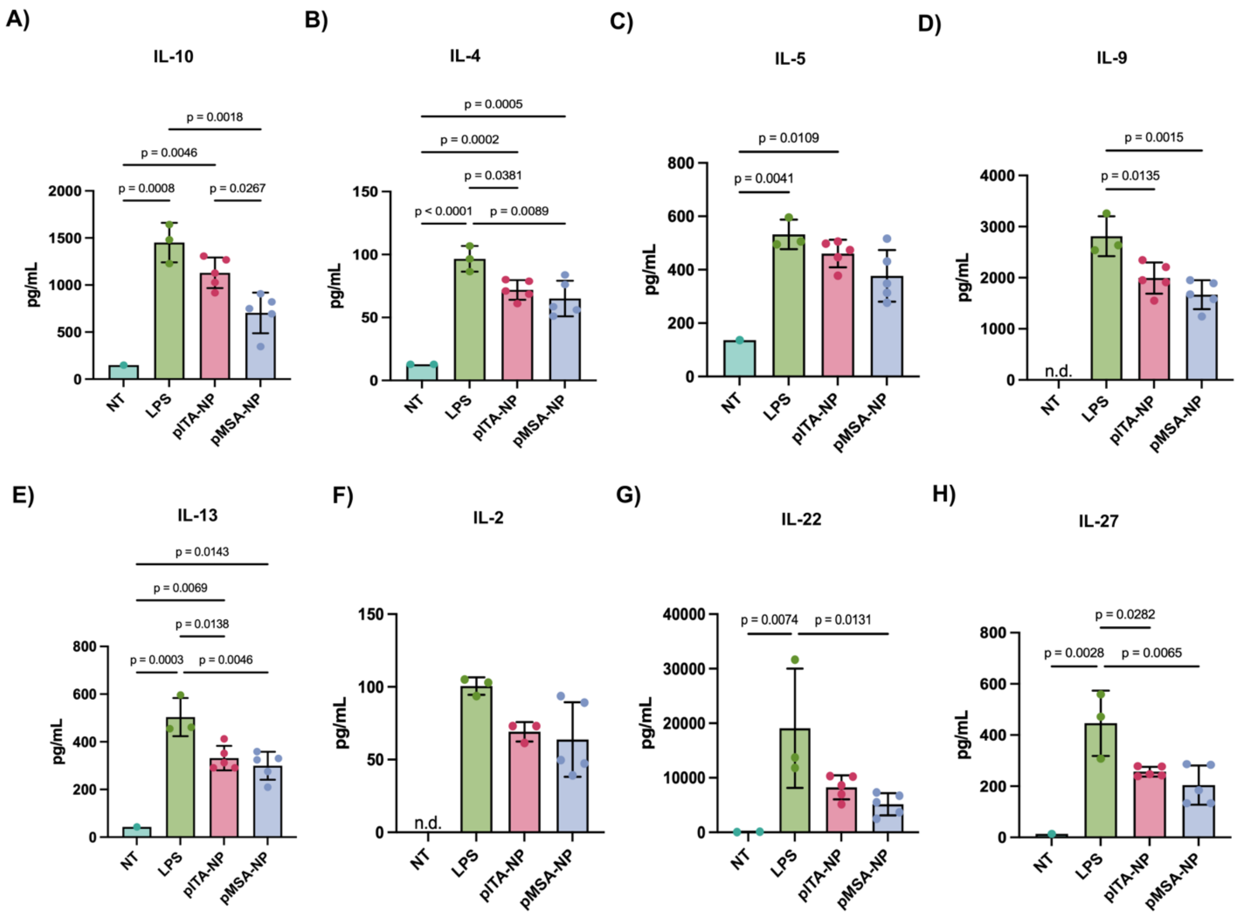


**Figure S15. Circulating regulatory and type 2-associated cytokine responses following metabolite-derived nanoparticle pretreatment in LPS-induced endotoxemia.** Concentrations of **(A)** IL-10, **(B)** IL-4, **(C)** IL-5, **(D)** IL-9, **(E)** IL-13, **(F)** IL-2, **(G)** IL-22, and **(H)** IL-27 in untreated (NT), LPS-treated, pITA-NP-treated, and pMSA-NP-treated mice. Data are presented as mean ± SD, with individual points representing individual animals. Statistical comparisons were performed using the Kruskal–Wallis test followed by Dunn's multiple-comparisons test; exact adjusted P values are shown for the indicated comparisons. Cytokine concentrations are reported in pg $mL^{-1}$.
